# Interplay of disordered and ordered regions of a human small heat shock protein yields an ensemble of “quasi-ordered” states

**DOI:** 10.1101/750646

**Authors:** Amanda F. Clouser, Hannah E.R. Baughman, Benjamin Basanta, Miklos Guttman, Abhinav Nath, Rachel E. Klevit

## Abstract

Small heat shock proteins (sHPSs) are nature’s “first responders” to cellular stress, interacting with affected proteins to prevent their aggregation. Little is known about sHSP structure beyond its structured α-crystallin domain (ACD), which is flanked by disordered regions. In the human sHSP HSPB1, the disordered N-terminal region (NTR) represents nearly 50% of the sequence. Here, we present a hybrid approach involving NMR, hydrogen-deuterium exchange mass spectrometry, and modeling to provide the first residue-level characterization of the NTR. The results support a model in which multiple grooves on the ACD interact with specific NTR regions, creating an ensemble of “quasi-ordered” NTR states that can give rise to the known heterogeneity and plasticity of HSPB1. Phosphorylation-dependent interactions inform a mechanism by which HSPB1 is activated under stress conditions. Additionally, we examine the effects of disease-associated NTR mutations on HSPB1 structure and dynamics, leveraging our emerging structural insights.

## INTRODUCTION

Small heat shock proteins (sHSPs) are a class of molecular chaperones that help maintain cellular proteostasis. Like other heat shock proteins, sHSPs are believed to interact with exposed hydrophobic regions of partly unfolded or misfolded proteins to help prevent irreversible aggregation, but unlike other heat shock proteins, they perform their functions independent of ATP. sHSPs are implicated in numerous human diseases on the basis of inherited mutations in the protein sequence or upregulation in certain cancers^1^. Cellular stressors such as oxidation and acidosis can influence their function^2–5^, and stress-induced phosphorylation of sHSPs typically increases their chaperone activity^6–8^. Despite their important roles in health and disease, relatively little is known about sHSP structure or structure-function relationships compared to other classes of chaperones.

Several properties make mammalian sHSPs particularly challenging for structural characterization. First and foremost, many sHSPs form large homo- and/or hetero-oligomers, and many of these exist as broad distributions of oligomeric species that contain different numbers of subunits. Furthermore, subunits exchange rapidly among oligomers, making it difficult to capture a single oligomeric state, or even a narrow distribution of states. Second, up to 50% of the sequence of sHSPs is believed to be intrinsically disordered and is unresolved in the few available structures of full-length sHSPs. Third, there is often local conformational heterogeneity even within ordered regions of the protein. Finally, the transient and promiscuous nature of interactions between sHSPs and client proteins makes it challenging to capture a sHSP/client complex of the sort that has fueled structural-mechanistic understanding of other chaperones. Our goal has been to develop hybrid strategies capable of providing structural/dynamical residue-level information on these challenging yet highly important systems.

sHSPs consist of three regions that also define different types of inter-subunit interactions in homo-oligomers (Fig. 1A). Most current structural information on human sHSPs is based on the structures of their defining feature, the central α-crystallin domain (ACD)^3, 9–13^. On their own, ACDs form dimers that are amenable to solution NMR and protein crystallography. Both sequentially and structurally conserved among the human paralogs, ACDs form an IgG-like β-sandwich fold of six or seven strands in truncated (ACD-only) structures and in the few oligomeric models and structures determined to date (Fig. 1B)^14–16^. ACDs dimerize by anti-parallel alignment of their β6+7 strands to form a long β-sheet dimer interface (ACD-ACD interaction). The flanking regions on either side of the ACD are far less conserved among paralogs and are predominantly disordered. The C-terminal region (CTR) is a relatively short extension that contains many charged residues and is highly flexible and disordered. CTRs are thought to serve as solubility tags for sHSPs. Most human sHSPs known to exist as large oligomers contain a three-residue motif of alternating Ile or Val residues known as the “IXI” motif near the beginning of their CTR. IXI motifs can bind in a “knob-and-hole” fashion into a groove formed between the top and bottom sheets of the ACD β-sandwich (the “β4/β8 groove”, Fig. 1B), defining a second type of intermolecular interaction observed in sHSP oligomers (ACD-CTR interaction). Finally, the N-terminal region (NTR) is a relatively long extension (50-100 residues in vertebrates) that is presumed to be disordered based on secondary structure prediction and the lack of density in electron microscopy (EM) and X-ray crystallography-based structures of sHSPs^17–19^. Relative to typical disordered regions, the NTR is enriched in hydrophobic and aromatic residues. Short regions of order have been observed in the NTR of HPSB5 by solid-state nuclear magnetic resonance (NMR)^14^ (Fig. 1 Supp1), in a crystal structure of HSPB6 in complex with a client protein^20^, and in a crystal structure of an HSPB2/HSPB3 heterotetramer^16^. The lack of sequence conservation in the NTRs among paralogs (Fig. 1C) raises the question of whether there are similarities among structural features of NTRs of other sHSPs or whether each protein is idiosyncratic. As the NTR drives sHSP oligomerization and is thought to play a critical role in its chaperone activity, the paucity of structural insights has greatly slowed progress towards understanding these important proteostasis components.

**Figure 1.**
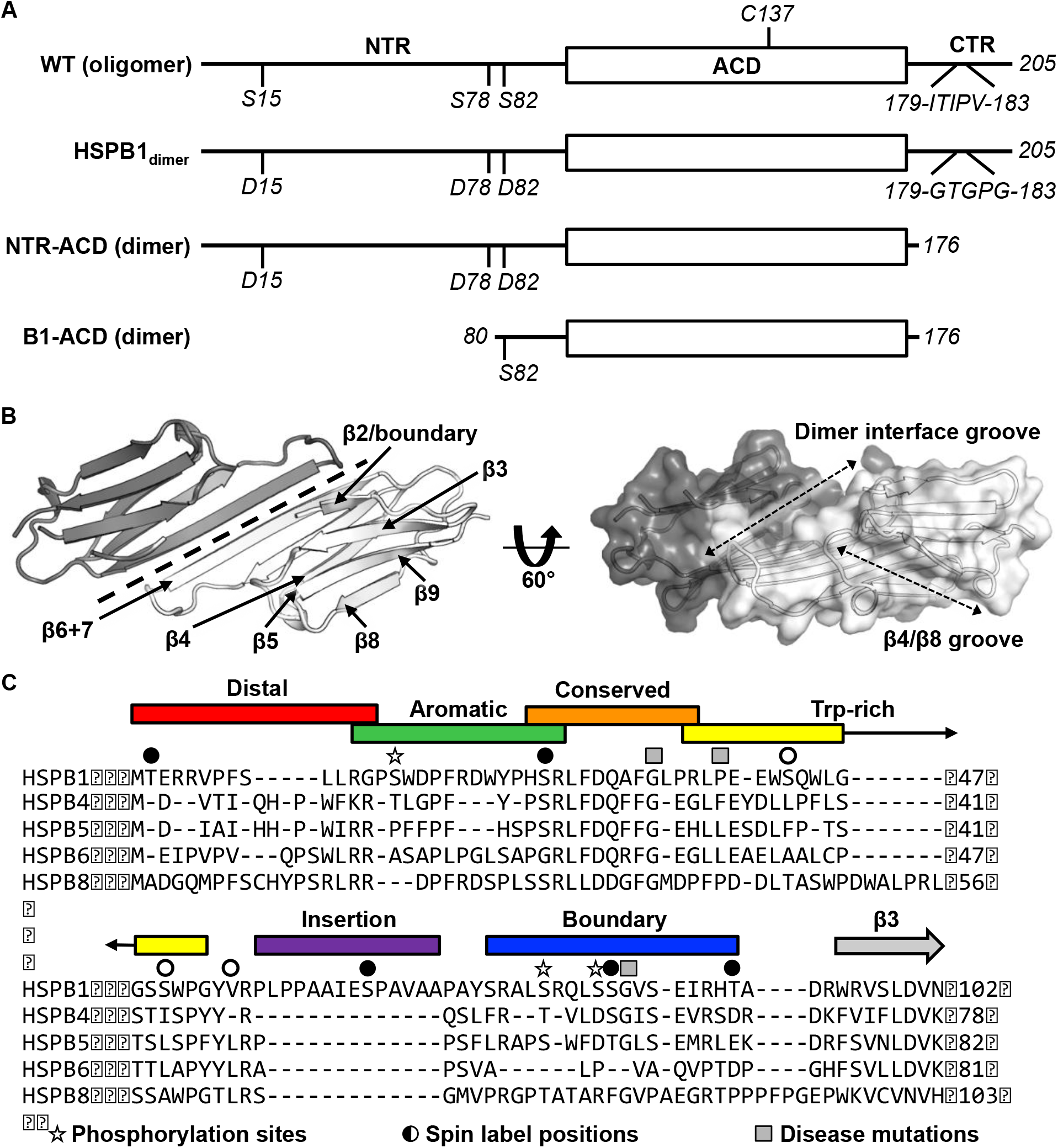
sHSP domain architecture and sequence alignment. (A) All sHSPs are defined by a conserved α-crystallin domain (ACD), that is flanked by N-terminal and C-terminal regions (NTR and CTR, respectively). Constructs used in this study are shown and their quaternary structure (oligomer vs. dimer) is indicated. HSPB1 contains a single cysteine, located in the ACD, which was substituted with serine in NMR-PRE and HDXMS experiments across several constructs. Phosphorylation-mimicking mutations in the NTR along with mutation of the IXI motif in the CTR (to abrogate ACD-CTR interactions) yields a full-length dimer, used in HDXMS experiments here. The C-terminally truncated construct (NTR-ACD) was used predominantly for NMR assignments and PRE experiments. The N-terminally and C-terminally truncated B1-ACD construct was used for peptide-binding NMR experiments. (B) ACD homodimer structure (4MJH), with dotted line indicating axis of symmetry along which ACDs interact via β6+7 strands. Dotted arrows in the right panel indicate the axes of the dimer interface and β4/β8 grooves. (C) NTR alignment of better-characterized human sHSPs shows minimal sequence conservation aside from the “conserved” region. The NTR sequence of HSPB1 was divided into six sub-regions for this study, which were probed in peptide form (with the exception of the highly hydrophobic insertion region). Sites of interest in this study are also indicated: 1) phosphorylation sites mutated to aspartate to mimic phosphorylation, 2) sites that were targeted for spin label attachment, and 3) three disease-associated mutations (G34R, P39L, and G84R).

Among the most ubiquitously expressed of the ten human sHSPs, HSPB1 is implicated in multiple biological roles in many cellular pathways, diseases, and tissues. In addition to its canonical chaperone role, HSPB1 has been implicated in the apoptosis pathway and interacts with cytoskeleton proteins^21, 22^. Upregulation of HSPB1 has been observed in certain types of cancer and has therefore drawn attention as a potential therapeutic target^23^. HSPB1 is phosphorylated within minutes after cells are subjected to stress by stress-related kinases at three NTR serine residues that are conserved among HSPB1 orthologs (Fig. 1A, C, and Supp1)^24^. Biochemical investigations have demonstrated that phosphorylated HSPB1 or phosphomimetic mutants form oligomers that are much smaller and more active than the distribution formed by unmodified HSPB1^25–29^. Several inherited mutations in HSPB1 are reported to be associated with the severe neurological disorders Charcot-Marie-Tooth disease and distal hereditary motor neuropathy. Disease-associated mutations are harbored in all three regions of the protein, with overrepresentation in the NTR and in residues near the dimer interface of the ACD and most appear to be autosomal dominant. Little is known about the structural mechanism for phosphorylation-dependent dispersion of HSPB1 into smaller oligomers or the structural effects of disease-associated mutations.

Here, we present the first residue-level structural analysis of full-length HSPB1, using a hybrid experimental approach of NMR spectroscopy and hydrogen-deuterium exchange mass spectrometry (HDXMS). Our study sheds light on the structural consequences of stress-induced phosphorylation and disease-associated mutations in HSPB1. We focus first on a dimeric species that represents a fully dissociated, stress-activated form of HSPB1 (“HSPB1_dimer_”) that is as effective at delaying the aggregation of the client protein, tau, as oligomeric wild-type (WT) HSPB1^30^. The dimer is generated by three phosphorylation-mimicking substitutions in the NTR (S15D, S78D, and S82D) and substitution of the CTR IXI motif to “GXG” (^179^ITIPV^183^ to GTGPG) (Fig 1A). Due to its monodisperse nature and relatively low molecular weight, the construct is amenable to residue-level analysis by solution-state NMR. Application of HDXMS to HSPB1_dimer_ revealed regions of the NTR that are protected from exchange, consistent with the presence of transient order. We defined sub-regions of the NTR based in part on HDXMS behavior, as illustrated in Figure 1C, and performed two types of NMR experiments. NMR binding experiments using peptides containing the NTR sub-regions and PRE experiments in which spin labels were placed in each sub-region. Together, these approaches revealed that, although the NTR is predominantly disordered, there are nevertheless specific interactions between the NTR and ACD in the context of full-length HSPB1. HDXMS data obtained for WT-HSPB1 oligomers show remarkably similar patterns of protection, consistent with the interactions and order defined in the HSPB1_dimer_ being conserved in the context of oligomeric HSPB1. Finally, HDXMS analysis of disease-associated mutations in HSPB1 reveals that the NTR is highly sensitive to single residue changes, resulting in non-local structural effects.

Altogether, our results show that even in a monodisperse form of HSPB1, there is substantial conformational heterogeneity, with multiple, specific contacts between regions of the NTR and the ACD. These contacts are altered in activation-mimicking and disease-associated mutated states, shedding light on the mechanisms by which perturbations such as phosphorylation or mutation can influence sHSP structure and function. The experimental approach presented here can be applied to other structurally heterogeneous systems that have proven difficult to study by traditional means, particularly those containing a mixture of ordered and disordered regions.

## RESULTS

### The “disordered” NTR makes extensive contacts with the ACD

Atomic-level structural information for HSPB1 ACD and CTR regions is available from crystallographic and NMR studies^11–13, 31^. Given the crucial yet enigmatic role of the disordered NTR in sHSP oligomerization and function, we sought to expand structural studies to include the NTR. Although oligomeric forms of HSPB1 are too large to analyze by traditional solution-state NMR and are too heterogeneous to crystallize, our previously reported HSPB1_dimer_ is amenable to solution-state NMR approaches^30^. A construct in which the CTR is truncated to the same position as the end of our B1-ACD construct (residue 176) is also dimeric in the phosphomimic context (termed “NTR-ACD”, Fig. 1A). Thus, HSPB1_dimer_ and its truncated form provide the first opportunity to obtain residue-level information of a sHSP with its NTR in solution. The simplest model for HSPB1_dimer_ would be a structured ACD dimer with flexible, disordered NTRs and CTRs that behave independently of other domains. Such a species would give rise to an NMR spectrum that would resemble that of the isolated ACD plus resonances that correspond to “random coil” positions. The NMR spectrum of an HSPB1 construct that lacks both its NTR and CTR (“B1-ACD”, Fig. 1A) and forms a well-structured dimer has been assigned^12^. Remarkably, few peaks overlap perfectly in overlays of ^1^H-^15^N HSQC spectra of B1-ACD and NTR-ACD (black versus blue, respectively; Fig. 2A), Therefore, the model in which the ACD behaves independently of its flanking domains is inaccurate, demanding investigation of interactions between domains.

**Figure 2.**
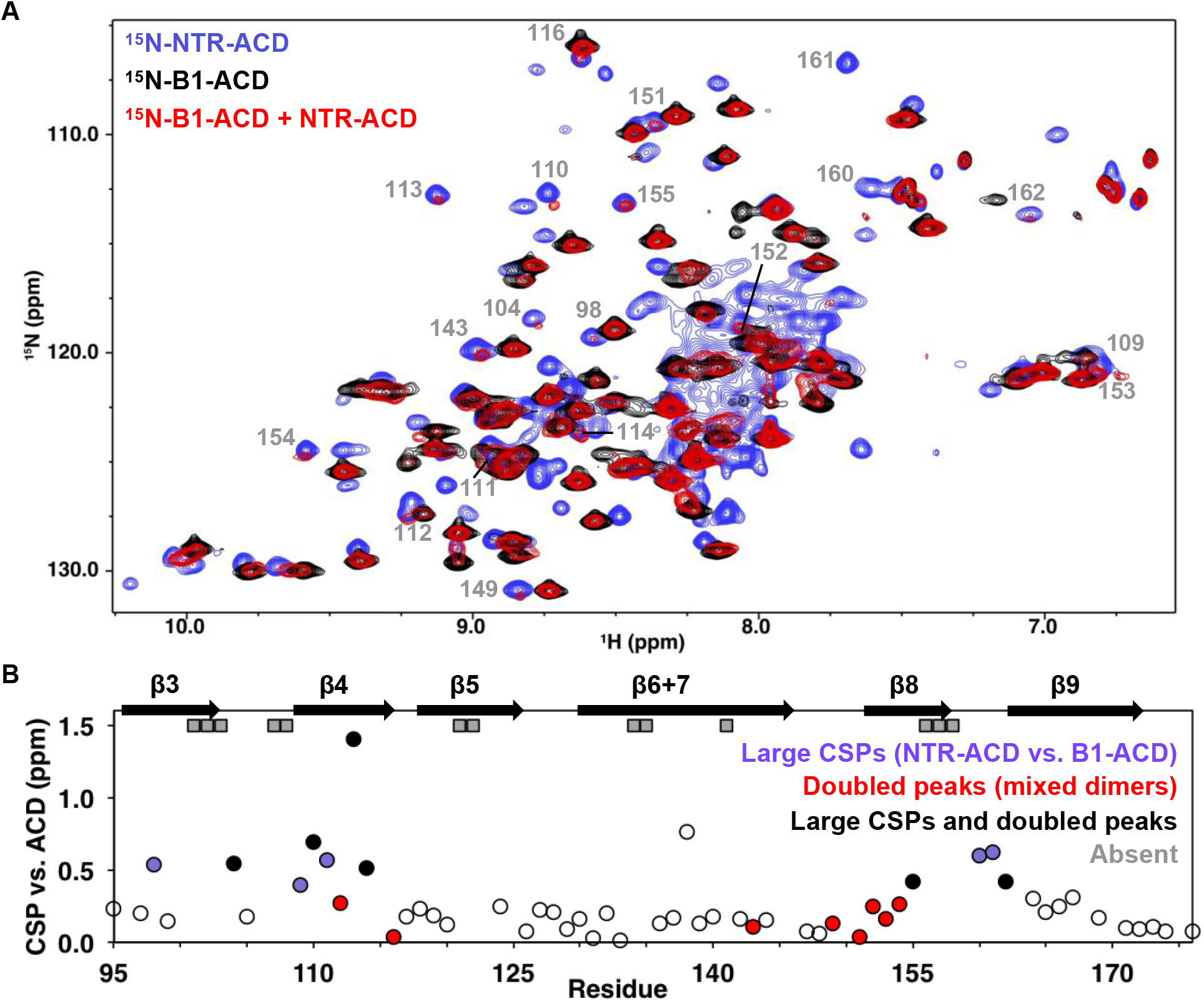
NMR analysis of NTR-ACD reveals changes in the ACD and increased heterogeneity when the NTR is present. (A) Comparison of B1-ACD (black), NTR-ACD (blue), and a mixture of ^15^N-B1-ACD and unlabeled NTR-ACD (red) ^1^H-^15^N HSQC-TROSY spectra. (B) CSPs measured for NTR-ACD compared to B1-ACD. The following color-coding highlights regions of interest: **blue**, residues most perturbed in NTR-ACD compared to ACD; **red**, residues that show NTR-ACD-like chemical shifts in the ACD mixing experiment, and **black**, residues that show effects for both cases. Gray squares correspond to residues in NTR-ACD whose resonances are missing and are presumably in substantially different chemical environments between B1-ACD and NTR-ACD.

The NMR spectrum of NTR-ACD overlays well with that of full-length HSPB1_dimer_, confirming that the CTR lacking its IXI motif does not interact detectably with the rest of the protein in the otherwise phosphomimetic context. (Fig. 2 Supp1). NTR-ACD was therefore used for subsequent NMR studies to limit spectral overlap of CTR peaks with NTR peaks in the central portion of the HSQC spectrum corresponding to disordered peaks. Most peaks corresponding to ACD residues could be identified and assigned from standard heteronuclear triple resonance NMR spectra collected on NTR-ACD (Fig. 2 Supp1, assignments deposited in BMRB). Despite the lack of precise overlap between the spectra of NTR-ACD and B1-ACD, the majority of residues in both contexts give rise to similar chemical shifts (^1^H, ^15^N, and ^13^C). The similarities of the chemical shift “fingerprints” indicate that the ACD structure is retained in the two contexts. Therefore, the widespread ^15^NH chemical shift perturbations (CSPs) observed for ACD resonances indicate differences in environment due to proximity of the NTR.

The largest perturbations in ACD peaks are for residues in the β3, β4, and β8 strands and loop L3/4 (Fig. 2B). These structural elements compose two grooves on the ACD dimer, known as the β4/β8 groove and the β3/β3 or “dimer interface” groove (Fig. 1B). In some cases, resonances appear to be absent in the NTR-ACD spectrum altogether: we were unable to identify peaks for several residues despite having assignments for these in the B1-ACD spectrum (gray squares in Fig. 2B). Analysis of 3D-heteronuclear spectra used to assign the NTR-ACD spectrum failed to identify peaks with similar ^13^C_α_/^13^C_β_ chemical shifts for these residues, implying that resonances for these residues are not observed and likely undergoing intermediate exchange between different conformations and/or chemical environments. As there is no evidence of slow chemical exchange in the context of B1-ACD, the broadening is likely due to dynamics and/or heterogeneity arising from the NTR. Altogether, the large perturbations indicate that the NTR interacts with the β4/β8 grooves at either end of the ACD dimer and the dimer interface at the center of the ACD.

Some ACD residues have multiple peaks in the NTR-ACD spectrum, indicating that they populate different chemical environments that interconvert slowly. The smallest difference in frequencies observed between multiple peaks from a single residue is 2 Hz, indicating a lifetime greater than 500 milliseconds. Notably, no peak doubling is observed in the B1-ACD spectrum. We could assign multiple peaks for four residues: positions 110 and 114 (flanking the β4 strand), position 148 (in loop L7/8, preceding the β8 strand), and residue 123 (in loop L5/6 between β5 and β6+7 strands. These positions are all near regions that exhibit CSPs and/or exchange broadening described above.

To investigate sources of the perturbations in ACD peaks in the NTR-ACD context, we examined NMR spectra of a mixture of ^15^N-B1-ACD and unlabeled NTR-ACD. Under reducing conditions (to inhibit disulfide bond formation at the dimer interface), mixed dimers composed of one ^15^N-B1-ACD subunit and one NTR-ACD subunit can form. This allowed us to observe perturbations in an ACD due to the NTR of its dimeric binding partner. As shown in Figure 2A, new peaks are observed in the spectrum of the mixture (red spectrum) that align with peaks in the NTR-ACD spectrum (blue spectrum). Peaks that exhibit this behavior nearly all correspond to residues in the β4/β8 groove and its flanking loops, predominantly the same residues whose peaks exhibit the largest CSPs compared to their position in the ACD-only spectrum (Fig. 2B). The congruence of certain peaks in the mixed-dimer spectrum with those in the NTR-ACD spectrum provides unambiguous evidence that their resonance positions arise from an interaction involving the NTR of the other subunit within a dimer. The identity of these peaks reveals that a site of interaction is the β4/β8 groove and its flanking loops. Additionally, many other peaks in the mixed-dimer spectrum lose intensity, shift, and/or change shape as compared to the B1-ACD spectrum. Such behavior is indicative of increased heterogeneity caused by multiple processes. Peaks that undergo such changes correspond to residues in the β3 strand and preceding residues and at the end of the β9 strand and start of the CTR (Fig 2 Supp2). Overall, the extent of perturbations observed in this mixing experiment reveals wide scale NTR-ACD contact and establishes that NTR/ACD interactions can occur between subunits within a dimer.

### Distinct regions of the NTR bind different ACD interfaces

To ask whether specific NTR regions interact with the ACD, we subdivided the NTR sequence into six sub-regions, referred to here as the distal region, aromatic region, conserved motif, tryptophan-rich region, insertion, and boundary region (Fig. 1C). We used NMR to test the ability of peptides from each region to bind to B1-ACD. As described below, peptides from the distal, aromatic, conserved motif, and boundary regions all caused distinct changes (either chemical shift perturbations, loss of peak intensity, or both) in the ^15^N-HSQC of the ACD, implying specific interactions of these regions with the ACD. No perturbations were observed for the Trp-rich peptide, providing confidence that the effects observed for other peptides are specific.

We also performed paramagnetic relaxation enhancement (PRE) experiments to detect interactions between the NTR and the ACD within the context of the NTR-ACD construct. PRE from a spin label broadens resonances of residues proximal to the label. The sole native cysteine at position 137 was substituted with serine and multiple constructs were created in which a single cysteine residue was introduced at NTR sites to which the spin label MTSL was conjugated. Spin label positions were selected to probe each NTR region, as shown in Figure 1C. Before performing NMR experiments, each Cys mutant was assessed for alterations in secondary structure and oligomeric properties (by CD and SEC, respectively, data not shown). Only variants that retain HSPB1_dimer_-like properties were investigated further. ^15^N and spin label were incorporated into these species and HSQC spectra of labeled dimers were collected with the active spin label and with the MSTL quenched by ascorbate. Quenched spectra were compared to the NTR-ACD spectrum to confirm that MTSL conjugation did not significantly perturb the constructs. PREs were quantified as ratios of peak intensity in active vs. quenched MTSL spectra. A range of behaviors were observed, depending on the spin label position: 1) strong, discrete effects, 2) smaller but still localized effects, or 3) widespread general effects.

As summarized in Figure 3, the results from the PRE experiments agree well with the peptide-binding studies, confirming that the interactions observed in the peptide-binding studies are recapitulated in the full-length protein. Results from individual peptides and spin-labels are described below and are provided in Figures 4, 5, and 4 Supp1.

**Figure 3.**
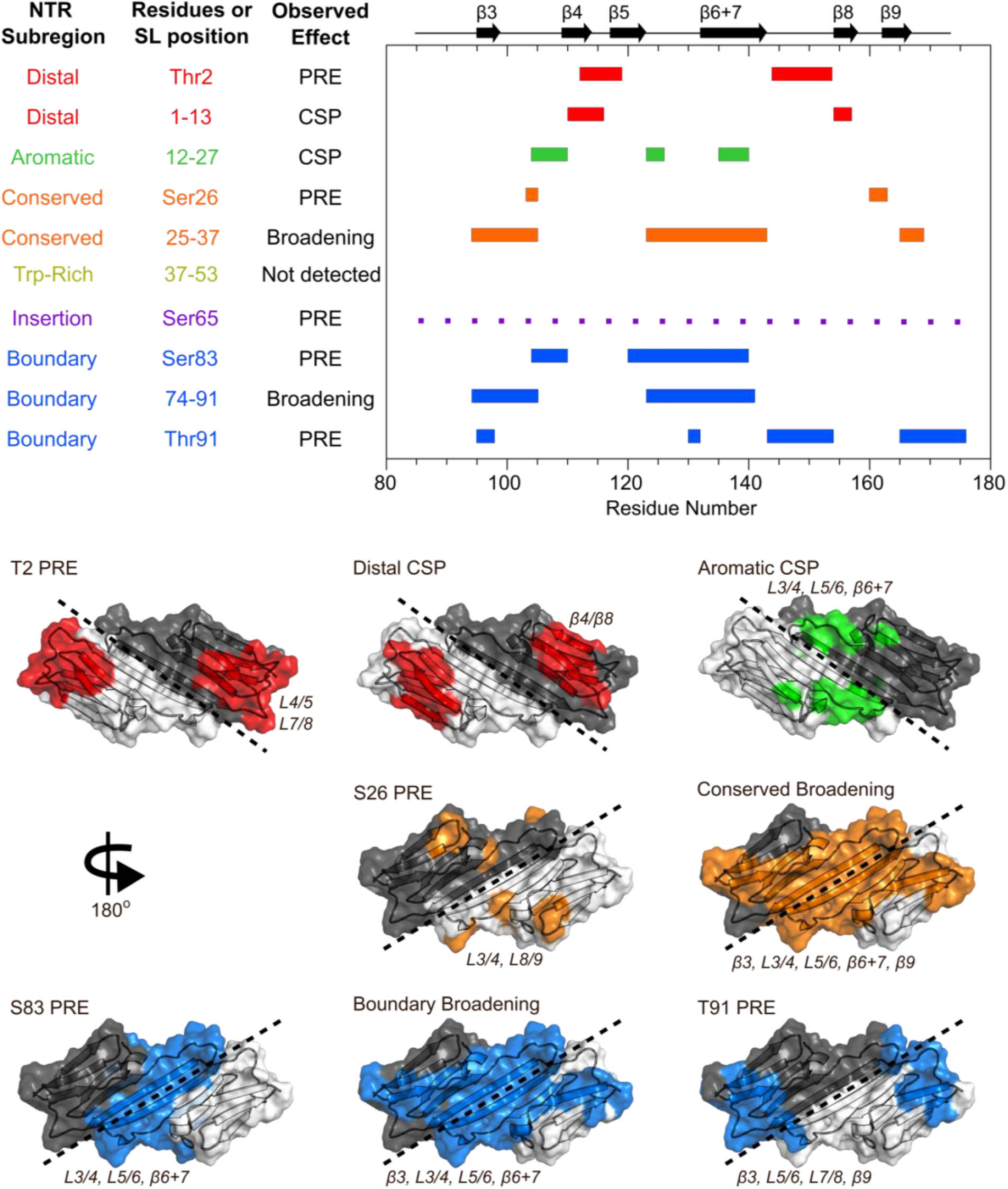
Summary of results from peptide-binding and PRE NMR experiments. Regions of the ACD that are perturbed upon peptide binding or lose intensity in the presence of a spin label are highlighted on the HSPB1 primary structure (top) and NMR structure (PDB 2N3J, bottom). The peptide from the Trp-rich region did not cause significant perturbations, and the presence of a spin label at position 65 caused widespread, nonspecific intensity loss, so these were not included in the bottom panel.

**Figure 4.**
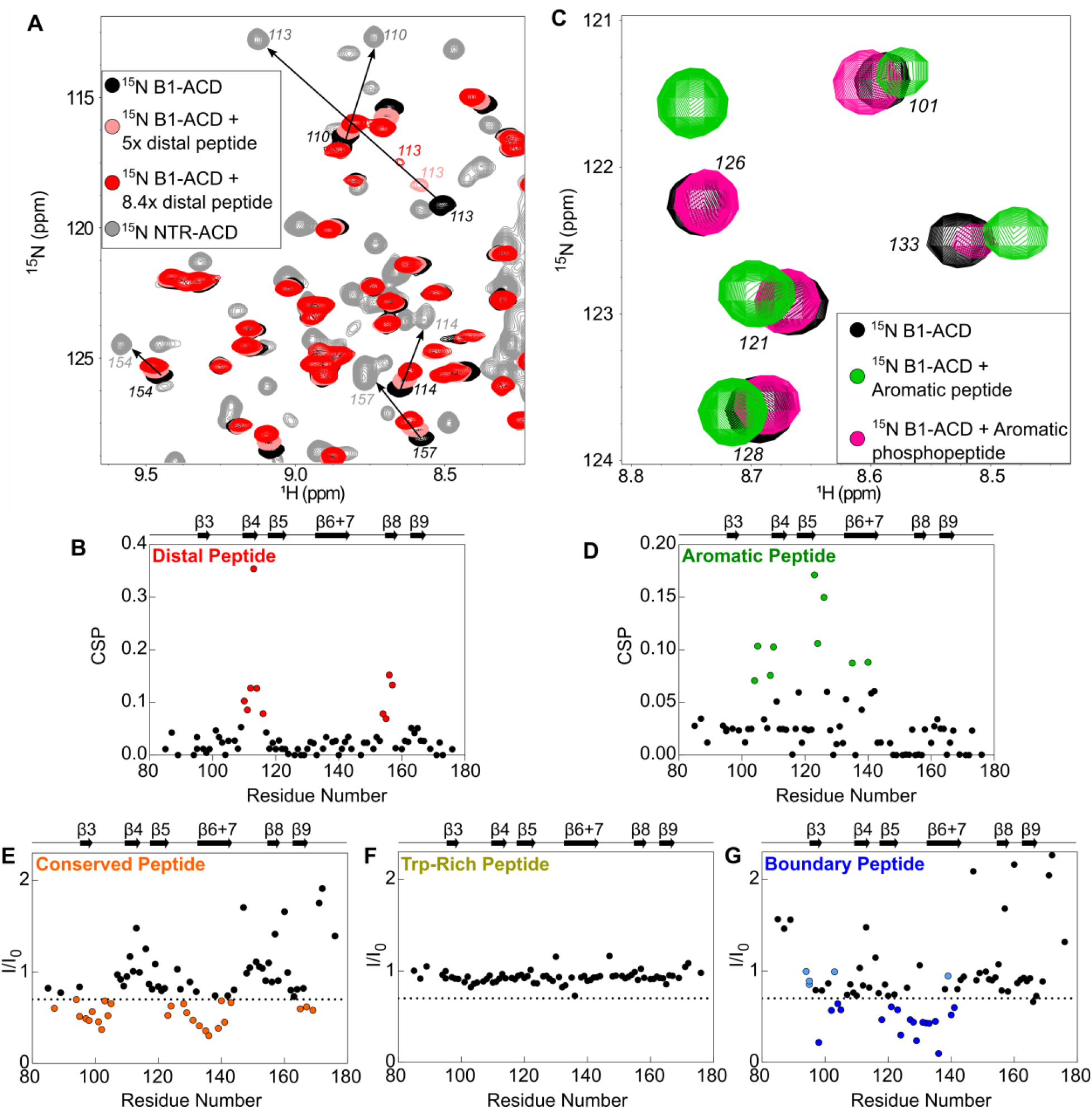
Perturbations of ^15^N-B1-ACD due to peptide binding. (A) The distal peptide, consisting of HSPB1 residues 1-13, causes CSPs in the ^15^N-HSQC spectrum of B1-ACD (black). Peak shifts (pink: 5 molar equivalents, red: 8.4 molar equivalents) occur along a trajectory toward the peak positions of the same residues in the ^15^N-HSQC of NTR-ACD (grey). (B) The strongest CSPs (red dots) map to residues in the β4-β8 groove. (C) The aromatic peptide (residues 12-27) causes CSPs in the spectrum of B1-ACD (green), but these are weakened when the peptide contains phosphoserine at site 15 (pink). (D) The CSPs map to residues in loops 3/4 and 5/6 and strand β6+7 (green dots). (E) The conserved peptide (residues 25-37) causes intensity loss in peaks in the ^15^N-HSQC spectrum of HSPB1 ACD corresponding to strands β3, β6+7, and β9. Peaks that lose more than 30% of their original intensity are colored in orange. (F) The Trp-rich peptide (residues 37-53) does not cause significant CSPs or intensity loss. (G) The boundary peptide (residues 74-91) causes both CSPs and intensity loss. Peaks that lose over 30% of their initial intensity are colored dark blue, and peaks with CSPs >0.05ppm that do not lose 30% of their initial intensity are colored light blue.

Addition of the distal peptide to ^15^N-B1-ACD caused distinct chemical shift perturbations (CSPs) in the ^15^N-HSQC spectrum that map to the β4-β8 groove (Fig. 3, 4A and 4B). Importantly, peaks shift along a trajectory between their positions in the spectra of B1-ACD and NTR-ACD (as indicated by the arrows in Fig. 4A). This is strong evidence that the large perturbations observed for these peaks in the NTR-ACD spectrum (relative to B1-ACD) are due to binding of the NTR distal region to the β4-β8 groove. The peptide binding was not saturated in the NMR experiments, thereby producing smaller CSPs relative to the NTR-ACD spectrum. The effective concentration of the peptide when attached to the ACD in its native form will be in the millimolar range, so the effects observed when the peptide is added in *cis* are relevant to the native state. The results add clarity to the ^15^N-B1-ACD/NTR-ACD mixing experiment described above (Fig. 2A). The new peaks that appear in positions that correspond to those observed in the NTR-ACD spectrum are due to distal region binding to the β4-β8 groove of the other subunit of the dimer in a “domain swap” relationship. We cannot rule out the possibility of a similar intra-chain interaction, but if it occurs it must be essentially identical to the inter-chain one observed in the mixed dimer.

The spectrum of NTR-ACD with MTSL at position 2 has strong peak intensity loss in two distinct sequence regions that correspond to loops L7/8 and L4/5, both of which lie near one entrance to the β4-β8 groove (Fig. 3 and 5A). Other peaks in the spectrum are largely unaffected by the spin label (i.e., I_para_/I_dia_ ∼1.0). This remarkably discrete PRE effect from a spin label at the extreme N-terminus of HSPB1 indicates that when the region is near the ACD, it inhabits a highly localized position. Furthermore, the PREs are consistent with only one of the two possible orientations of distal region binding in the groove, namely aligned parallel to the β8 strand and antiparallel to β4 strand such that residue 2 only contacts residues near the beginning of β8 and the end of β4. This is the opposite orientation from that observed for the CTR IXI/ β4-β8 groove binding observed in a crystal structure of HSPB1-ACD (4MJH)^13^. Distal region binding to the β4-β8 groove has been observed in crystals of HSPB6^20^ and an HSPB2/3^16^ complex, but those sHSPs contain canonical IXI motifs in their NTR. HSPB1 does not contain such a motif in its distal region; we propose instead that alternating hydrophobic residues in the HSPB1 segment ^6^VPFSLL^11^ bind, supporting the idea that other hydrophobic amino acids can participate in β4-β8 binding. Our results thus identify a novel interaction and indicate that motifs from both the NTR and CTR of HSPB1 can bind in the β4-β8 groove but are oriented in opposite directions within the groove.

**Figure 5.**
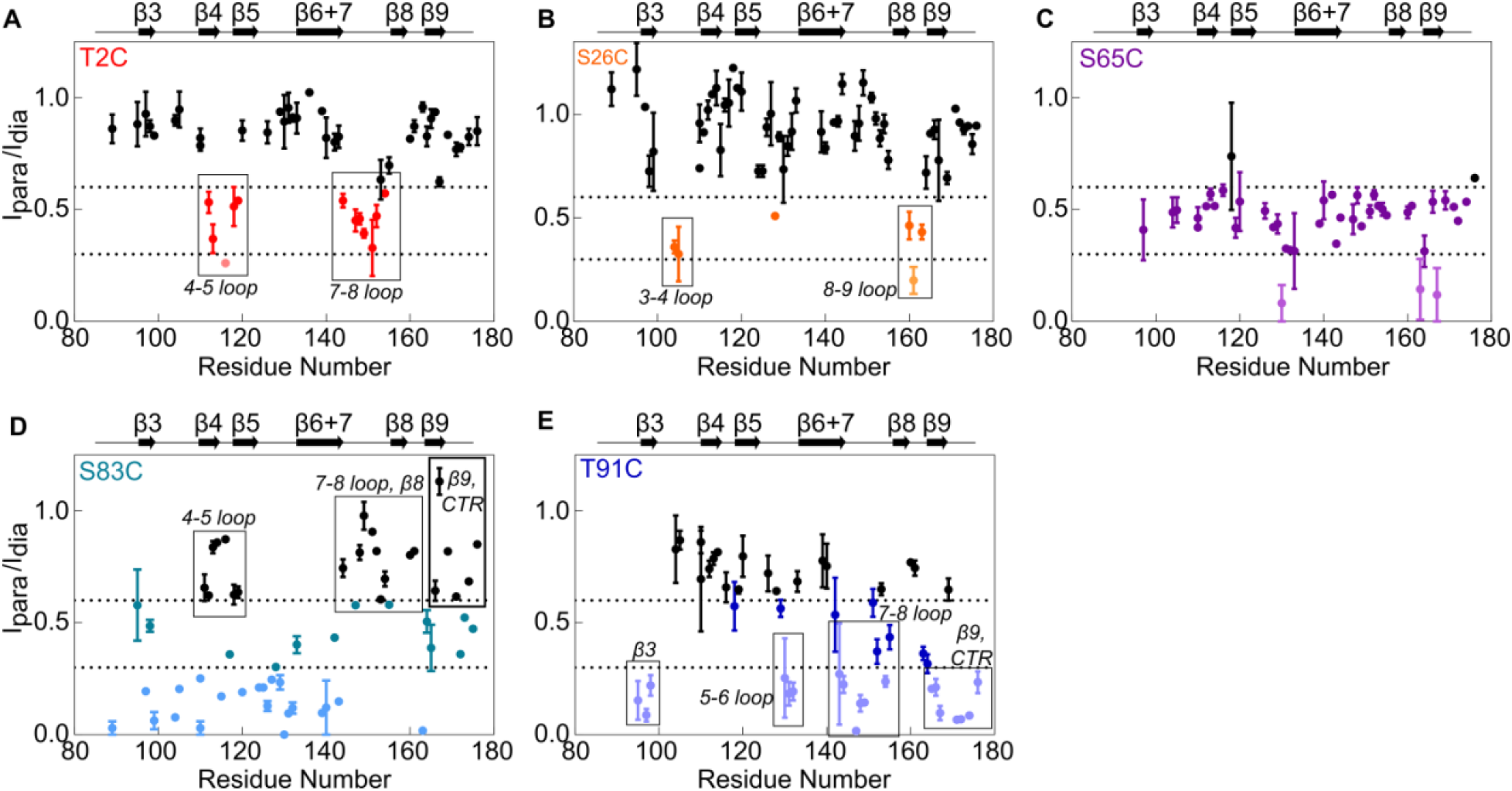
Paramagnetic relaxation enhancement NMR reveals contacts between NTR sites and residues in the ACD. The spin label MTSL was conjugated at five positions in the NTR of ^15^N-NTR-ACD, and spectra were collected in the presence of the active spin label (paramagnetic) and after it had been quenched by ascorbate (diamagnetic). Error bars represent the range of two independent experiments. Peaks that lose more than 40% of their intensity are highlighted in color, and peaks that lose more than 70% are shown in lighter colors.

Peptide binding and PRE results indicate that the conserved motif binds at the ACD dimer interface groove (i.e. strands β3 and β6/7 from each subunit), and that the aromatic sub-region spans between the β4-β8 groove and the dimer interface, connecting the distal region to the conserved motif. The aromatic peptide (residues 12-27) caused distinct chemical shift perturbations in peaks corresponding to one side of the ACD, primarily loops L3/4 and L5/6, as well as some residues in β6+7 (Fig. 3, 4C and 4D). This is consistent with the orientation inferred for the distal region, and shows that as the NTR exits the β4-β8 groove, it contacts the side of the ACD via loops L3/4 and L5/6 (Fig. 3). The aromatic region contains one of the three HSPB1 phosphorylation sites (Ser15, Fig. 1C), so we also tested binding by a version of the aromatic peptide containing phospho-serine at position 15. For the most part, the CSPs are absent or markedly reduced (Fig. 4 Supp1A), indicating that phosphorylation of Ser15 serves to release the aromatic region from the ACD. We note that both L3/4 and L5/6 are enriched in negatively-charged amino acids (^100^DVNHFAPDE and ^124^HEERQDEHG, respectively), suggesting an electrostatic mechanism by which phosphorylation at Ser15 could regulate HSPB1 structure and activity.

Conserved motif binding at the ACD dimer interface is indicated by intensity loss in peaks of residues in β3, β6+7, and in loop L3/4 upon addition of the conserved peptide to ^15^N-B1-ACD (Fig. 3 and 4E). A spin label at the aromatic region/conserved motif boundary (position 26) causes subtle peak intensity loss localized to loops L3/4 and L8/9 leading out of the interface groove, as would be expected if the conserved motif is bound in the groove and the aromatic region runs along one side of the ACD between the β4-β8 groove and the dimer interface groove (Fig. 5B). Conserved motif binding at the dimer interface groove has been observed in HSPB6 and in HSPB2/3 hetero-tetramer crystals^16, 20^. Given the high conservation of this motif and of residues that compose the dimer interface, it is likely that a similar relationship exists in HSPB1.

Although the Trp-rich peptide showed no detectable interactions with B1-ACD (Fig. 4F), we were able to obtain some insights regarding its behavior. While attempting to introduce spin labels, we found the Trp-rich region to be highly sensitive to mutagenesis and were unable to probe its interactions through PRE experiments. Cysteine substitutions of S43 and V55 within the NTR-ACD construct yielded species larger than dimers and not amenable to NMR. An S50C NTR-ACD construct was dimeric but conjugation of MTSL at this position produced larger species. While the altered oligomeric propensity rendered these mutants unsuitable for PRE experiments, the ability of changes in this region to override the dimer-promoting mutations implicates the Trp-rich region in maintaining the delicate balance among oligomeric species. This sub-region harbors the disease-associated mutation P39L, which also affects HSPB1 oligomeric properties (see later section).

The insertion region was probed by a spin label at position 65, near the center of the region. Intermediate intensity loss was observed in most B1-ACD peaks, with some stronger effects in β9. The widespread but modest PREs suggest that the insertion region does not make sustained contact with any specific region of the ACD but rather “hovers” over all faces of the ACD. (Fig. 5C) The region was omitted from peptide binding experiments due to its hydrophobicity and insolubility in aqueous buffer. Nevertheless, we were able to obtain backbone assignments for this region, which gave further insight into its behavior (discussed in next section).

Finally, the boundary region appears to interact at the dimer interface groove, as evidenced both by peptide binding and PRE results (Fig 3, 4G, 5D, and 5E). Addition of the boundary region peptide (residues 74-91) to B1-ACD caused intensity loss and CSPs in peaks corresponding to residues near the dimer interface (Fig. 3 and 4G). We were able to incorporate spin labels at two boundary region positions, 83 and 91, which flank the predicted β2 strand (residues 86-88 in PDB 4MJH). The spin label at position 91 (S91-SL) caused distinct intensity loss in peaks from loops L5/6 and L7/8 and from residues at the beginning of β3 and end of β9 (Figure 3 and 5E). The highly localized PREs on one side of the ACD are congruent with residues 91 being near the beginning of the β3 strand. In contrast, the spin label at position 83 caused wide-spread intensity loss in peaks corresponding to the dimer interface and the opposite side of the ACD (Fig 3 and 5D). In this case, the ACD residues that are *not* affected by the spin-label are more informative: the data clearly show that position 83 does not approach loops L4/5, L7/8, and L9/CTR, all of which are localized to one side of the ACD dimer (the side affected by S91-SL). The fact that spin labels at positions flanking the putative β2 strand hit opposite sides of the ACD provides strong evidence that the boundary region spans the dimer interface groove in an antiparallel direction to strand β3, consistent with formation of a β2 strand. Whether this interaction is mutually exclusive with binding of the conserved motif to the dimer interface or can occur simultaneously cannot be deciphered from these experiments.

A boundary-region peptide that contains phospho-serine residues at positions 78 and 82 caused nearly identical perturbations to the non-phosphorylated peptide (Fig. 4 Supp 1B), indicating that phosphorylation neither directly disrupts nor enhances the interaction between the boundary region and the dimer interface groove. However, the PRE results from S83-SL indicate that this region approaches residues that are perturbed by the aromatic peptide (Fig. 3 and 5D), suggesting that the boundary region phosphorylation sites may be close in space to the S15 phosphorylation site, despite being over 60 residues apart in sequence.

### Conformational heterogeneity is observed throughout the NTR

Peaks that are not assigned to ACD residues in the ^1^H-^15^N HSQC spectrum of HSPB1 presumably arise mainly from the NTR. Many of these peaks are weak (but reproducible across multiple protein preps) and were not observed in low-sensitivity 3D heteronuclear spectra despite high levels of deuteration and use of high field magnets equipped with cryoprobes. For resonances that were detected in heteronuclear spectra, there was considerable degeneracy in chemical shifts (C_α_/C_β_), likely due to intrinsic disorder and/or conformational heterogeneity. While these properties hampered unambiguous assignment of NTR peaks, the weak and/or poorly-resolved peaks indicate that a majority of residues in the NTR exist in multiple environments, leading to their peak broadening. There are two clear exceptions: we were able to assign a contiguous stretch of residues that span the insertion and boundary regions and several residues in the Trp-rich region.

Peaks corresponding to residues 62-79 could be assigned (Fig. 6A); these span most of the (non-proline) residues of the insertion region and the start of the boundary region (Fig. 1C). All peaks assigned for this stretch have high intensities (similar to CTR, Fig. 6C) and chemical shifts consistent with random coil structure (Figure 6A). Only the N-terminal part of the boundary region is assigned, while the latter part of the boundary region that is predicted to form β2 could not be assigned. Notably, residues 64-69 were each assigned to two distinct peak sets, indicating two conformational states that interconvert slowly, despite their both having “random coil” chemical shifts. Based on the observed chemical shift differences and the slow exchange condition, we can estimate an upper limit for the exchange rate of ∼ 25 s^-1^. The second conformation represents a substantial fraction of the population, as judged from relative peak intensities (>20%). A short contiguous stretch at the C-terminal end of the Trp-rich region (residues 46-49) was assigned, with Ser49 having two peak sets. Thus, both regions exist in at least two distinct conformations and/or environments. The observed behavior may arise from *cis* and *trans* conformations of one or more of the numerous proline residues in the insertion and Trp-rich regions. However, ^13^C chemical shifts needed to confirm proline isomerization could not be obtained from the data and we did not pursue mutational analysis to identify the source(s) of heterogeneity due to the abundant prolines in the regions.

**Figure 6.**
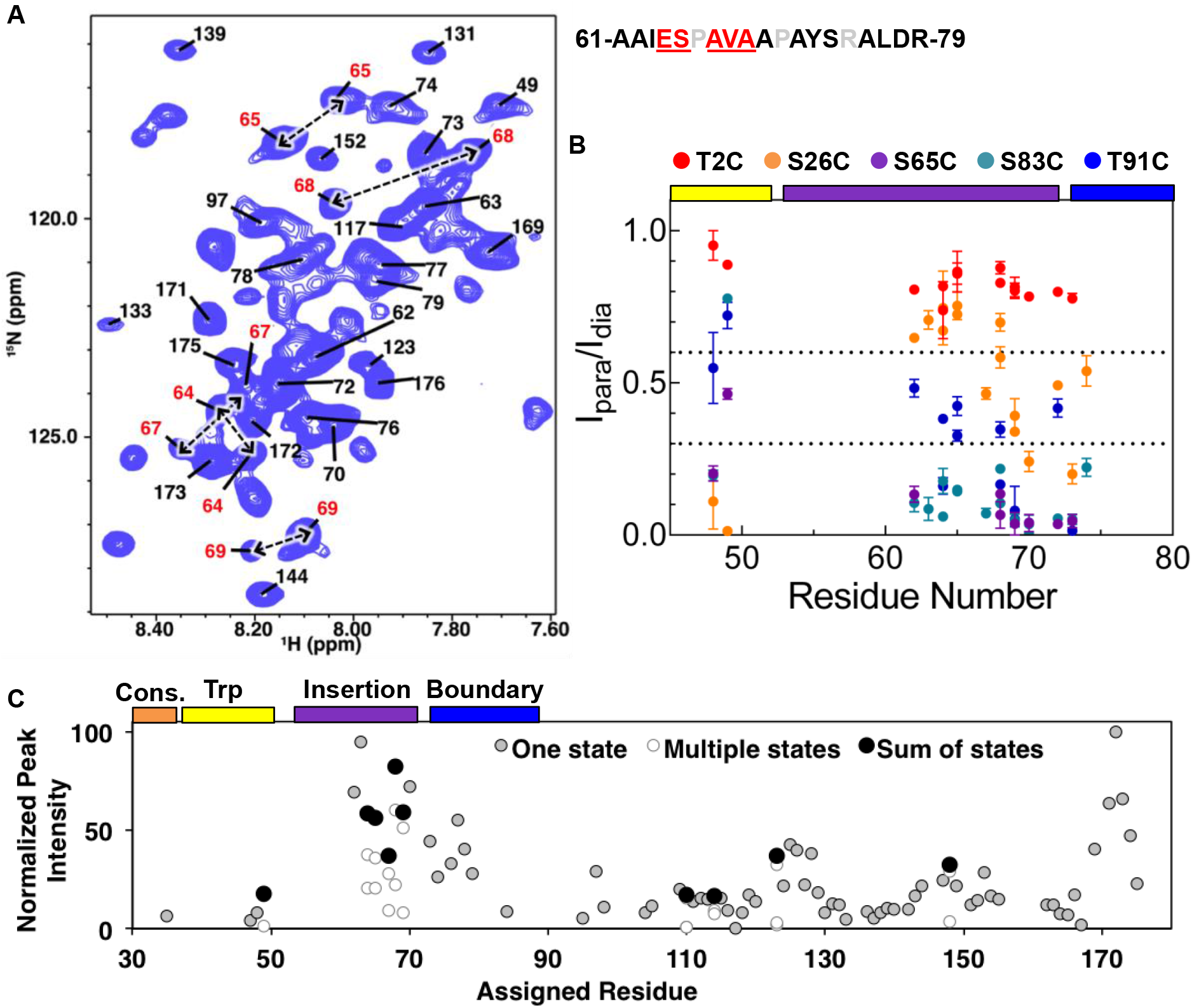
Assignment of NTR residues reveal disorder and heterogeneity. (A) ^1^H-^15^N HSQC-TROSY spectra of NTR-ACD, highlighting strong peaks in the center of the spectrum that correspond to disordered regions of the protein (NTR and CTR). For part of the NTR, pairs of peaks are assigned to the same residue indicating conformational heterogeneity (red in sequence and peak labels). (B) Summary of PRE effects from previously described spin label positions on assigned regions of the NTR. (C) Relative (most intense peak = 100) intensities of all non-overlapping peaks assigned in NTR-ACD, including sums of peaks corresponding to different conformations.

Assignments of NTR HSQC peaks provided information regarding proximities of NTR regions to each other from PREs (Fig. 6B). The spin label at insertion region residue 65 yields strong PREs across the assigned NTR peaks, validating their assignments to this region of sequence. The position 2 spin label did not yield PREs in assigned NTR residues or in weak unassigned peaks, indicating that the distal region’s locations do not overlap with those of the Trp-rich or insertion regions. A position 26 spin label at the beginning of the conserved region gives modest-to-strong PREs to the Trp-rich and boundary regions but weak PREs to the insertion region. The position 83 spin label gave strong PREs to the boundary and insertion regions, and the position 91 spin label gave moderate PREs to the same regions. Together, this provides evidence for extensive interactions between the sub-regions of the NTR. In particular, the conserved and Trp-rich regions seem to be in close contact with each other, as are the insertion and boundary regions. Only the distal region appears not to make extensive contact with other parts of the NTR. The data support a model in which the NTR exists in a compact state with extensive NTR/ACD and NTR/NTR contacts as well as some residual structure, rather than a more flexible, extended conformation seen for many IDRs.

We used HDXMS to compare NTR properties in dimeric and oligomeric forms of HSPB1. Backbone amide protons in structured and/or buried regions exchange with deuterons more slowly (i.e., are “protected” from exchange) than those in unstructured or accessible regions. Since HDXMS can be performed on proteins of any size under a variety of solution conditions it provided an opportunity to obtain peptide-level information on the dynamics of HSPB1 oligomers. Time courses of deuterium exchange from three seconds to one hour were measured at room temperature for C137S-HSPB1 oligomers and C137S-HSPB1_dimer_. The sole Cys residue, C137, that resides at the dimer interface was substituted to abrogate the need for reducing agent to inhibit disulfide bond formation across the dimer interface that could confound the analysis. Pepsin digestion yielded peptides across the full length of the protein (peptide statistics are shown in Fig. 7 Supp Table 2).

**Figure 7.**
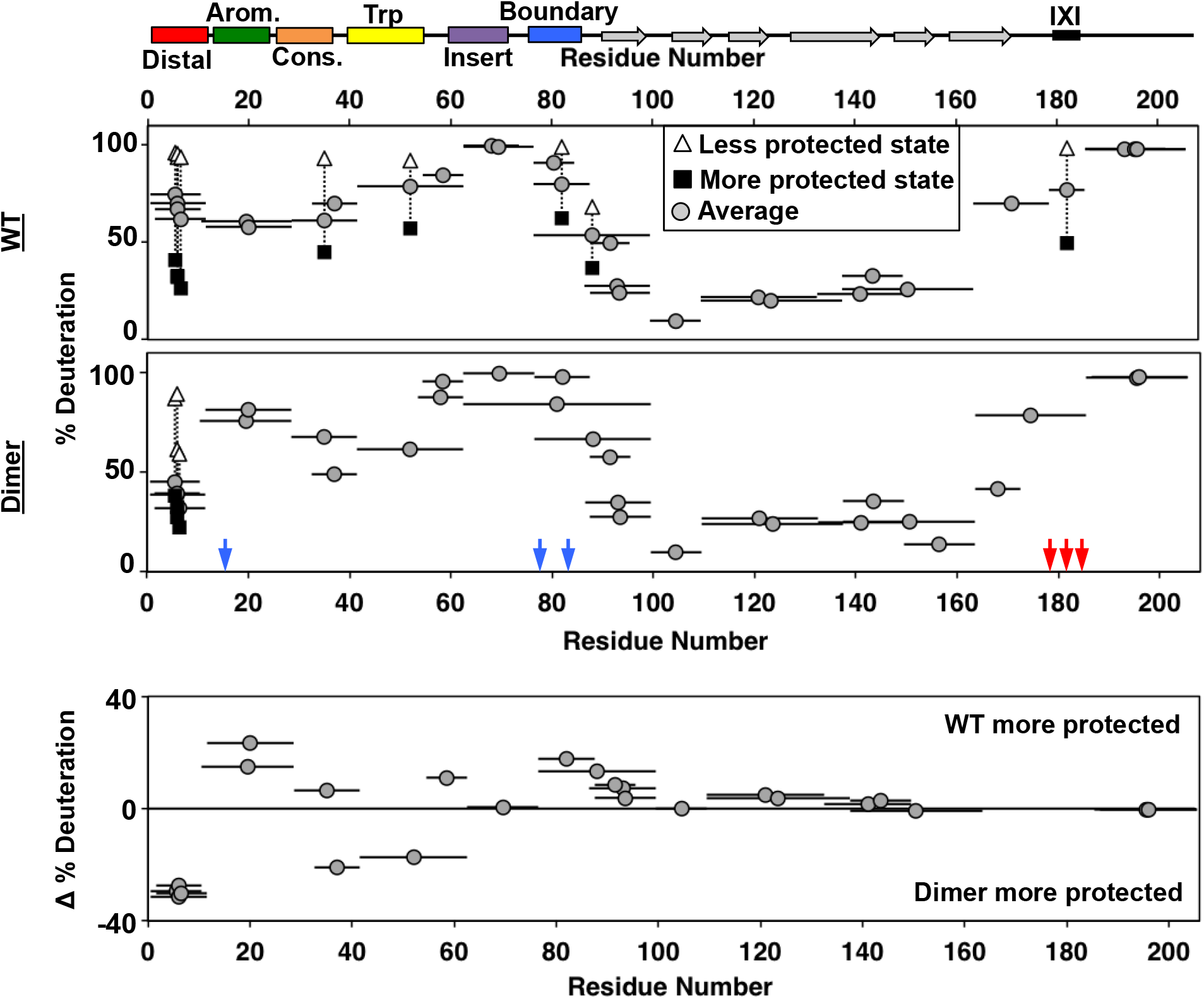
HDXMS analysis of WT oligomers and HSPB1_dimer_ reveals changes in protection and heterogeneity in the NTR. Representative peptides are indicated as horizontal bars. The midpoint of each peptide is represented by a gray circle showing the deuteration level of the peptide after 3 seconds (see Supplement for full kinetic table). For peptides that show a bimodal distribution (different states), black squares and white triangles represent the more- and less-deuterated populations, with the gray circle representing the weighted average deuteration. Blue and red arrows indicate sites of mutation used to generate HSPB1_dimer_. The difference in deuteration level between WT and HSPB1_dimer_ is shown for each peptide in the last plot. Although identical peptides for the start of the CTR cannot be compared due to the mutations introduced, the profile for the 164-185 peptide in HSPB1_dimer_ is consistent with the high level of deuteration observed for peptides 164-178 and 179-185 in WT oligomers.

Deuterium uptake levels differed most dramatically across regions of HSPB1 in the earliest (3 sec) timepoint, shown in Figure 7 (see Fig. 7 SuppTable1 for full kinetic information). The profile for HSPB1_dimer_ (middle panel, Fig. 7) is congruent with observations and conclusions drawn from NMR presented above. Residues 87-172, which span the ACD, show protection even at the latest time point, which is consistent with its β-sandwich structure. Peptides within the aromatic, insert, and boundary regions in the NTR, along with the far C-terminal region (residues 187-205) of the CTR are all highly deuterated within 3 seconds, consistent with them lacking stable secondary structure. Interestingly, peptides along the distal, conserved and Trp-rich regions of the NTR showed moderate protection, indicating the presence of some local structure. All peptides covering the distal region displayed a bimodal isotopic mass envelope at the early time points indicating that there are two distinct populations of the distal region: one that exchanges readily with deuterium and one that is protected (see Fig. S7 Supp1). The bimodal distributions observed for these peptides may reflect the populations of the NTR that are either bound and sequestered or free and solvent-accessible. A substantial protected population is present through 15 seconds, implying that the lifetime of the bound state is on the order of at least several seconds. All of the other peptides outside of the distal region displayed only unimodal spectra in the HSPB1_dimer_ dataset.

The general features of the exchange profile for oligomeric HSPB1 (top panel, Fig. 7) are similar to HSPB1_dimer_, with the ACD being most protected and the NTR and CTR less protected. As in the dimer, the insertion region, boundary region, and CTR have the highest rate of deuterium uptake, indicating that they remain unstructured and accessible in oligomers. However, the aromatic region shows more protection in oligomers than in HSPB1_dimer_, indicating some involvement in the oligomerization. The most striking feature of the oligomeric HSPB1 HDX profile is the large number of bimodal peptides that are observed. The distal region showed similar bimodal spectra as seen with the HSPB1_dimer_, except the relative population of the more protected species was diminished (∼40% vs. 80%). Beyond the distal peptides, another six peptides have two distinguishable populations in the context of HSPB1 oligomers. Peptides that arise from the conserved, Trp-rich, and boundary regions in the NTR display two populations. The conserved and boundary regions were observed to interact with the ACD in the dimer context from NMR results, so either these interactions are longer-lived within the confines of oligomers, or they may also be directly involved in oligomeric interactions. The presence of a substantial protected population at 4 minutes indicates a lifetime of interaction of several minutes in the oligomeric context. In the CTR, a peptide that contains the IXI motif (mutated to GXG in HSPB1_dimer_ to inhibit its binding to the β4/β8 groove) has a population (∼45%) that is protected from deuterium exchange and one that is completely exchanged. This behavior is consistent with the notion that IXI-containing CTRs exist in both free and β4/β8-bound states.

The populations observed for the bimodal spectra are not sufficiently resolved along multiple timepoints to determine precise rates of slow conformational exchange (“EX1 kinetics”) or to make quantitative comparisons. Nevertheless, qualitative comparisons at the well-resolved 3 sec timepoint can be made between regions or between protein constructs. The weighted average deuteration for each peptide with bimodal exchange in Figure 7 provides a qualitative estimate of the relative populations of “bound” and “free” states. For example, both the distal and IXI-CTR peptides show a substantial proportion (∼40 - 45%) of the region in a protected state. These two regions bind to the same ACD groove, creating a situation where there are twice as many potential β4/β8 groove-binding sequences as there are grooves. The HDX data indicate that enough groove-binding sequences are solvent-protected to imply that the majority of β4/β8 grooves are occupied in HSPB1 oligomers and that, on average, half the distal sub-regions and half the CTR-IXIs are bound.

To get a more complete overall picture of what else is different in the oligomeric form, we compared the deuteration levels for each identical peptide that could be identified between HSPB1_dimer_ and HSPB1 oligomers, using the weighted average percent deuteration for bimodal peptides. In the bottom panel of Figure 7, peptides that appear above the 0% line are more protected in oligomers and peptides that fall below the 0% lines are more protected in dimers. Overall, the ACD displays similar deuterium uptake regardless of whether it is in an isolated dimer or a large oligomer. This indicates that the extent of deuterium exchange at this timepoint is predominantly dictated by the β-sandwich structure. All peptides arising from the distal region show more protection in HSPB1_dimer_, with the weighted deuteration skewed to the more protected state (larger proportion) consistent with a larger fraction of the N-terminus bound in the dimer. This may be due to the CTR mutations introduced to generate the dimer, as the lack of IXI motif in the CTR removes it from competition for the β4/β8 groove. Two other contiguous peptides (33-41 and 42-62) that span conserved and Trp-rich regions show greater protection in the dimer. The overlapping 29-41 peptide is more protected in the oligomer, indicating that protection occurs in the first few residues of the peptide in oligomers, whereas protection occurs in later residues in the dimer, consistent with distinct interactions or structural features. The NMR results identified an interaction between the conserved region and the ACD dimer interface, but provided no evidence for an interaction involving the Trp-rich region and the ACD. According to secondary structure predictions, the Trp-rich region has the highest propensity of any NTR region to adopt secondary structure. Intriguingly, the CD spectrum of HSPB1_dimer_, but not HSPB1 oligomer, has an unusual positive peak at 230 nm^30^; similar features have been attributed to exciton couplets that can arise from interactions between Trp rings. Together, the HDX and CD data suggest that the Trp-rich region adopts specific structure in dimeric species that is not populated to a detectable degree in the oligomer. The two regions that are less protected in HSPB1_dimer_ are the aromatic and boundary regions, which contain the phosphorylation-mimicking mutations. The NMR data indicate that the conserved and boundary regions both interact with the dimer interface groove. The increased accessibility of the aromatic region is consistent with the reduced binding of the phosphorylated form of the aromatic peptide. It is possible that release of the aromatic region, which neighbors the conserved region, is coupled to decreased protection of the boundary region. Formation of a new structural feature involving the Trp-rich region might only be possible when neighboring regions are released from their ACD contacts.

### Modeling inter-region interactions in HSPB1_dimer_

The results presented above indicate that the HSPB1 NTR contains distinct regions that reside in specific locations relative to the well-defined ACD dimer and that, in many cases make direct contact with the ACD. We sought to combine the information garnered from the peptide-binding, PRE, and HDX experiments into a set of structural models. Our goals for this modeling process were two-fold. First, the models aid in visualization of the NTR-ACD interactions described above. Second, the modeling process allows us to determine which combinations of NTR-ACD interactions can generate physically realistic structural models, and thus whether any combination of NTR-ACD interactions may not be physically possible. These structures are intended neither as a complete sampling of HSPB1 conformational space nor as atomic-level models of specific interactions.

We produced a homology model by combining the crystal structure of the HSPB1 ACD (4MJH) with peptides from the crystal structure of the HSPB2/3 hetero-tetramer using the PyMOL Molecular Graphics System. The starting model included all of the possible NTR-ACD interactions identified above: two copies of the β2 strand, a distal motif bound in each β4/β8 groove, and a copy of the conserved motif bound in the dimer interface groove. Clashes between the HSPB1 ACD structure and the peptides from the HSPB2/3 crystal structure were eliminated and missing loops were modeled in using PyRosetta. In total, there are four possible ways to connect the fragments in the homology model: either copy of the distal motif in the β4-β8 grooves could be connected to the conserved motif at the dimer interface, and the conserved motif could be connected to either copy of the β2 strand (Fig. 8A). In total, we created four ensembles of 100 structural models, which each sampled a different way of connecting the NTR peptide fragments in our initial homology model (Fig. 8B, 8C, and 8 Supp1). We found that it was possible to develop realistic structural models that contain all types of NTR-ACD interactions identified above, as well as all possible loop connectivities. The dimer interface groove is able to accommodate two copies of the β2 strand in addition to the conserved motif, and both β4/β8 grooves are able to bind a copy of the distal motif. These peptide fragments can be connected to each other in all four conceivable ways.

**Figure 8.**
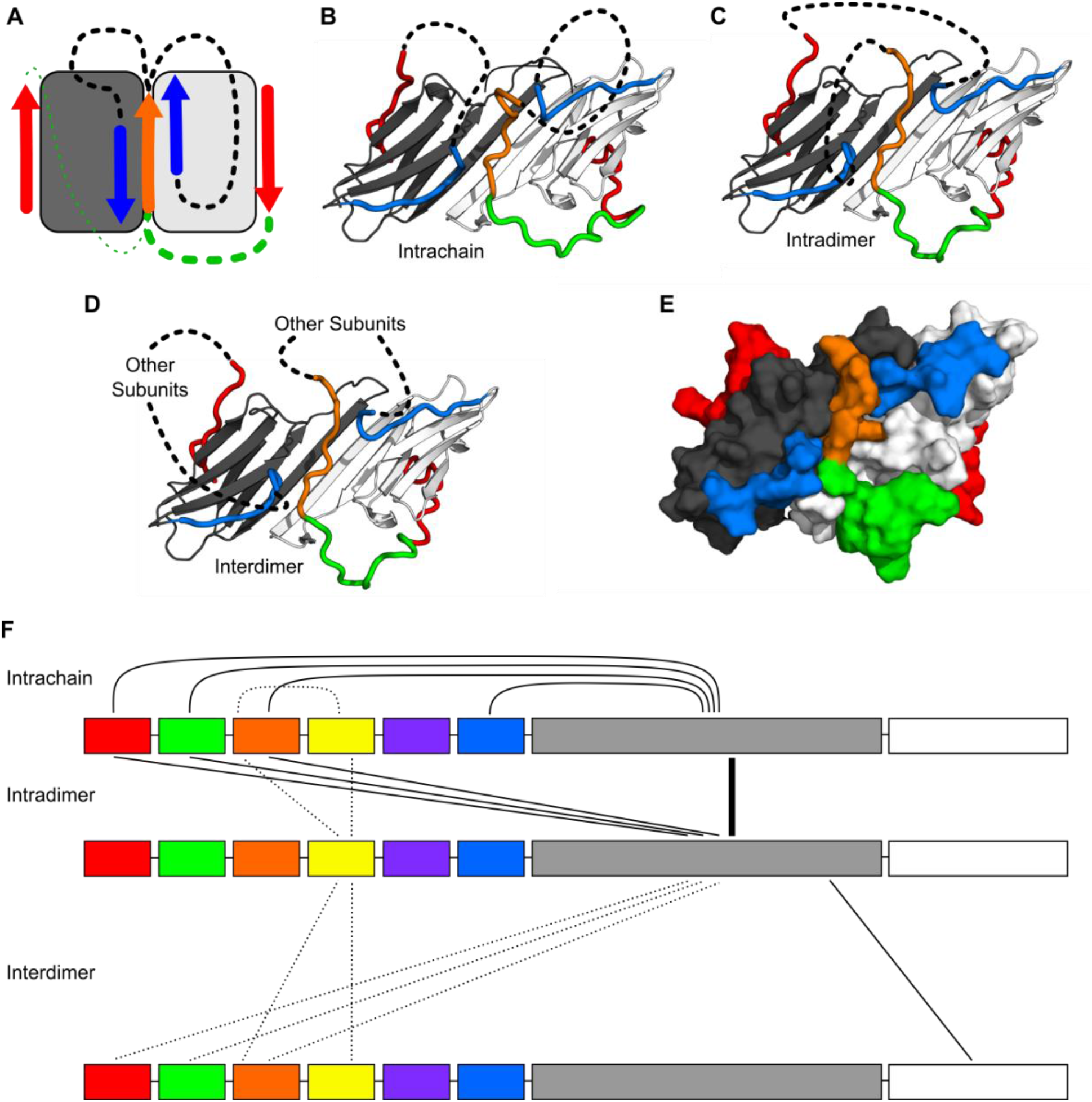
Modeling of NTR-ACD interactions. (A) Cartoon showing the starting structure of the ACD with two copies of the distal motif (red arrows), one copy of the conserved motif (orange arrow), and two copies of β2 (blue arrows). The missing loops can be modeled with four potential connectivities. Either copy of the distal motif can be connected to the conserved motif (green lines), which can then be connected to either copy of β2 (black lines). Based on our NMR results, we believe it more likely that the conserved motif is connected to the distal motif oriented in the opposite direction (thicker green line), so we only include structures with this connection in this figure. (B) If the conserved motif is connected to the β2 strand oriented antiparallel to it, the contacts between the distal and aromatic regions and the ACD occur within the same polypeptide chain. (C) If the conserved motif is connected to the other β2 strand, the contacts occur between different chains but within the same dimer. (D) It is likely that within the context of a higher-order oligomer, similar contacts could occur between subunits that are not part of the same ACD dimer building block. (E) A surface representation of the model in panel C shows that the regions of the NTR included in our model make extensive contact with the ACD and match the perturbed regions highlighted in Figure 3. (F) The types of intra- and interchain contacts that are possible within HSPB1 dimers and higher-order oligomers are outlined. Solid lines represent interactions for which we have evidence from our NMR, HDX, and modeling data. Dotted lines represent hypothetical interactions for which we do not have direct evidence but which we believe are likely to occur.

If the conserved motif is connected to the distal motif oriented in the opposite direction, the aromatic region forms a loop along the side of the ACD containing loops 3/4, 5/6, and 8/9 to connect these regions (Fig. 8B and 8C). Given our peptide-binding NMR results for the aromatic region, we believe that this configuration is favored, particularly in the non-phosphorylated state. Nevertheless, it is also physically possible for the aromatic region to connect the conserved motif to the distal region in the opposite groove, in which case it spans across the top of the ACD β-sandwich, contacting the β8, β9, and β3 strands (Fig. 8 Supp1). While we do not observe experimental evidence for this interaction, we cannot rule it out, particularly in the phosphorylated state in which the aromatic region has low affinity for the ACD. In either case, the conserved motif can then be connected to either one of the β2 strands, while the distal motif not connected to the conserved motif can be connected to the other. These connections consist of ∼50-75 residues and contain part of the boundary region, the insertion region, the Trp-rich region, and (for the chain with an unbound conserved motif) the conserved and aromatic regions. Given their length, they can adopt multiple conformations and orientations relative to the ACD. We have omitted these regions from the models shown in Fig. 8 and its supplement for clarity and to avoid over-interpretation of the structural aspects of regions for which we have limited experimental data. Surface representations of these models show that the locations of the NTR sub-regions are in good agreement with the ACD surfaces perturbed in the peptide binding and PRE NMR experiments (comparing Fig. 3 with Fig. 8E).

Overall, the results from the modeling suggest that any combination of the NTR-ACD interactions defined in our study is theoretically possible. While we only created models containing the maximum NTR-ACD interactions supported by our experimental data, any of the interacting motifs we have modeled could dissociate from the ACD and adopt a more disordered conformation. The results from our NMR and HDX experiments indicate that most of these NTR regions occupy both ACD-bound and ACD-unbound conformations, so it is likely that multiple combinations of NTR/ACD interactions occur in solution. Additionally, the similarity of protected regions in the HDXMS profiles of HSPB1 dimers and oligomers indicate that the interactions depicted in these models also occur within higher-order oligomers. The peptide fragments depicted in our dimeric models could conceivably be connected to other ACD dimers or monomers within an oligomer (Fig. 8D). The array of possible interactions within sHSP oligomers is depicted in Fig. 8F. Many regions can form intra-chain, intra-dimer, and inter-dimer interactions. The possibility for multiple combinations of interactions and connectivities contributes to the high degree of plasticity and heterogeneity observed for HSPB1 in NMR and HDX experiments.

### Disease mutants in the NTR have differential effects on HSPB1 structure

To leverage new insights regarding local structure in the NTR and interactions with the ACD, we sought to assess how reported disease-associated mutations in the HSPB1 NTR affect structure and/or dynamics. We chose two NTR disease mutants whose effects on oligomer size (larger than WT) and chaperone function have been previously characterized: G34R and P39L mutants in the conserved and Trp-rich regions, respectively^32^. To identify localized effects of the mutations, we performed B1-ACD-peptide binding experiments with mutant peptides. To detect potential global effects, we performed HDXMS analysis on mutant proteins. Glycine at position 34 is the final conserved residue in the “conserved motif” found among orthologs and paralogs; its mutation to Arg is expected to reduce flexibility of the backbone and/or alter the region’s interactions. At the start of the Trp-rich region, Pro39 is conserved among orthologs but not among paralogs. Notably, Pro39 is two residues before a region that is predicted to have helical propensity (residues 41-46, Fig. 1 Supp1). Mutation of Pro39 to leucine might change the structural propensity of this region by increasing flexibility and/or favoring helical formation. As presented below, we found highly divergent local structural effects of the two mutations, despite their close proximity in sequence.

As presented above, the conserved region peptide (residues 25-37) causes peak broadening in dimer interface groove residues (i.e., β3 and β6/7) in HSPB1 ^15^N-B1-ACD spectra. An otherwise identical peptide that contains the G34R substitution yields greatly reduced perturbations (Fig. 9B), indicating that the bulky, charged Arg sidechain disrupts the ability of the conserved region to interact with the dimer interface groove. Consistent with the notion that the environment of the mutated conserved region is altered, increased deuterium uptake for the fragment that contains the G34R substitution in otherwise wild-type HSPB1 (oligomer) is observed by HDXMS (Fig. 9A, bottom panel). The increase in deuterium uptake was sufficiently large (as high as 25% among the early time points), that it likely reflects a true increase in local flexibility, rather than a consequence of the G to R substitution altering the intrinsic exchange rate of this peptide^33^ Unexpectedly, the NTR boundary region also shows enhanced deuterium uptake in G34R-HSPB1 and, strikingly, no longer shows bimodal behavior. All other portions of oligomeric HSPB1 are not significantly changed in their exchange profiles.

**Figure 9.**
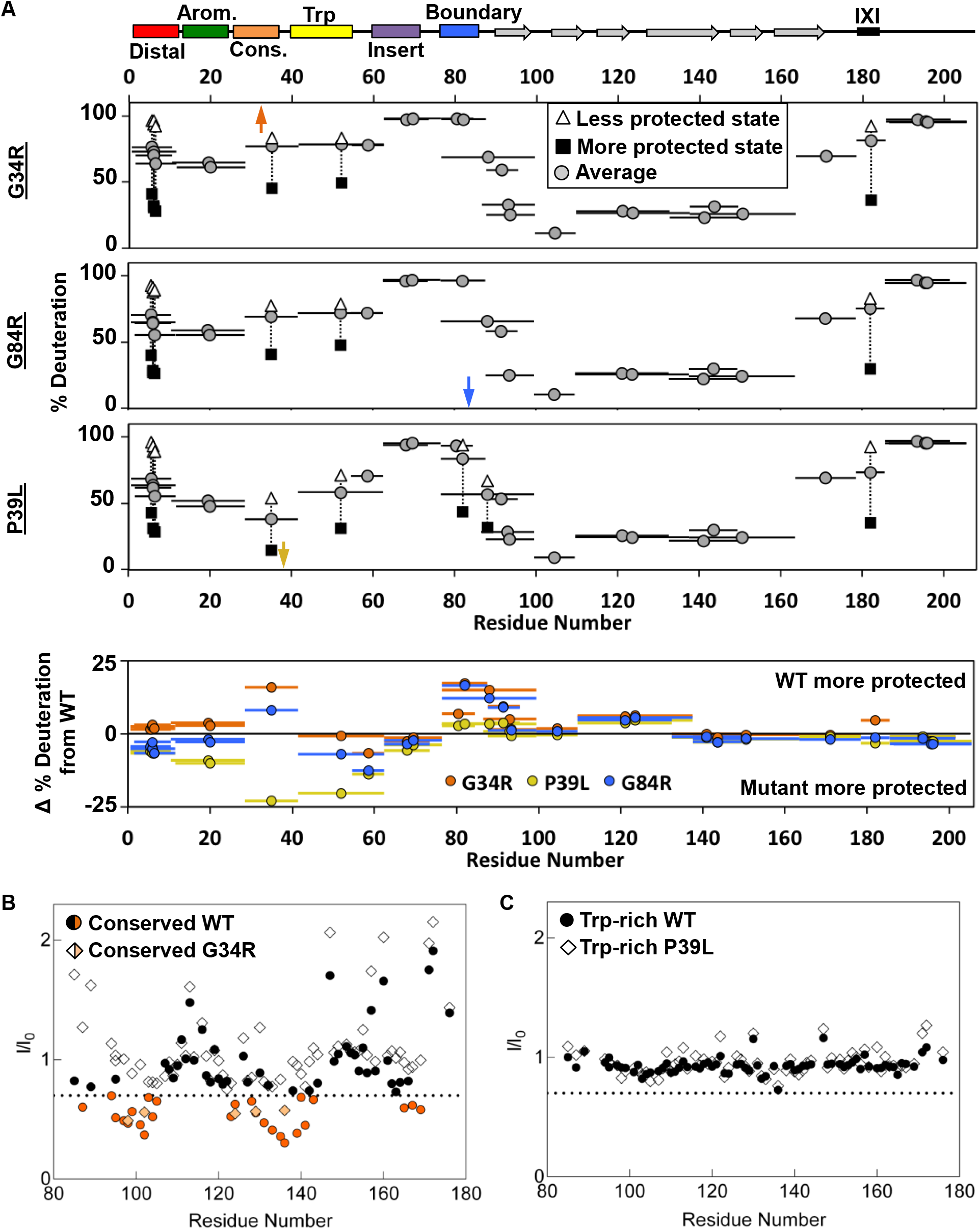
Analysis of disease-associated HSPB1 NTR mutations G34R, G84R, and P39L. (A) HDXMS analysis of each mutant. Representative peptides are indicated as horizontal bars. The midpoint of each peptide is represented by a gray circle showing the deuteration level of the peptide after 3 seconds (see Supplement for full kinetic table). For peptides that show a bimodal distribution (different states), black squares and white triangles represent the more or less deuterated populations, with the gray circle representing the weighted average deuteration. Arrows indicate sites of mutation (G34R, G84R, and P39L). The deuteration difference from WT for each disease mutant is shown for each peptide in the last plot. (B) NMR intensity loss in G34R-modified conserved peptide binding experiment with ACD. “WT” conserved peptide intensity ratios are shown as circles, and G34R peptide ratios as diamonds. Affected residues below the dotted line are colored orange. Fewer residues are affected by mutant peptide binding and to a lesser extent. (C) Analogous peptide binding experiment with Trp-rich peptide and P39L-modified peptide. In both cases, no notable intensity losses occur.

The non-local effect of the G34R mutation on the boundary region suggested to us that there could be coupling between the two NTR regions, similar to possible coupling between phosphorylation sites in the aromatic and boundary regions mentioned earlier. We took advantage of another identified disease-associated mutation that similarly substitutes arginine for glycine, but in the boundary region (G84R) to test this hypothesis. Intriguingly, HDXMS data on the mutant (Fig. 9A) revealed enhanced deuterium uptake in both the boundary region and in the conserved region, confirming that the disposition of these two non-contiguous NTR regions are inter-dependent. The NMR data indicate that both regions interact with the ACD dimer interface-groove, and our modeling suggests that both interactions can occur simultaneously. While the NMR data do not address whether the binding of one influences the other, the HDXMS data provide some clarity on this question. The observed inter-dependence of protection from deuterium exchange implies that each region enhances the ability of the other to contact the groove and that the conformation that leads to protection in the HDXMS experiment has both a conserved region and a boundary region present in the groove. Whether the two bound sub-regions are on the same HSPB1 polypeptide or different ones within a dimer or oligomer cannot be ascertained from these experiments, but our modeling implies that both cases are likely to occur.

As mentioned above, we did not detect evidence of binding for the Trp-rich region peptide to ^15^N-B1-ACD. We obtained similar results for a peptide that contains the P39L substitution, indicating that the mutation does not lead to a gain of binding function (Fig. 9C). Nevertheless, introduction of P39L into otherwise wild-type HSPB1 has a marked effect on the HDXMS profile (Fig. 9A). The first four NTR sub-regions are substantially more protected from deuteration in mutant oligomers, while the ACD is essentially unaffected. The largest increase in protection is observed in the fragment that contains the mutation (29-41). This comparison to WT is complicated by the addition of an extra amide from the P39L mutation, but the massive decrease in exchange (25%) is consistent with the predicted increased helical propensity in this region and an observed increase in helicity revealed in the CD spectrum (Fig. 9 Supp1). Altogether, the observations suggest that the P39L mutation increases local secondary structure and/or promotes NTR-NTR contacts. In further support of this conclusion, the subunit exchange rate for P39L-HSPB1 is three-fold slower than for either wild-type or G34R oligomers (Fig. 9 Supp1).

Altogether, the results highlight a variety of alterations that occur in the HSPB1 NTR under situations such as phosphorylation or disease mutation. These alterations can have profound non-local effects on other sub-regions of the NTR, leading to global changes in HSPB1 structure and oligomerization.

## DISCUSSION

Despite being among the most ubiquitously expressed of the ten human small heat shock proteins, there is a paucity of structural information regarding HSPB1. As for all known examples, the central ACDs of two subunits of HSPB1 adopt a dimeric β-sandwich structure^3, 11–13, 16, 20^. However, there has been a complete lack of structural information for the remaining ∼50% of HSPB1, most of which is represented by the enigmatic NTR. To overcome the challenges posed by high heterogeneity and polydispersity of large HSPB1 oligomers, we sought to obtain new structural information from a more tractable dimeric form of HSPB1 that retains its chaperone activity. Despite its monodispersity in solution, HSPB1_dimer_ exhibits substantial heterogeneity as detected by both NMR and HDXMS. Exemplary of the heterogeneity and dynamics, a majority of the NMR resonances from NTR residues are either of low intensity or are broadened beyond detection even at high magnetic fields, signifying multiple environments for these residues. The disordered NTRs of sHSPs have remained enigmatic for decades, with very little structural information emerging. The hydrophobic NTRs are required for oligomerization of some sHSPs (HSPB1, HSPB4, HSPB5), although paradoxically others remain predominantly dimeric despite having similarly long and hydrophobic NTRs (HSPB6 and HSPB8). Our effort to generate a well-behaved dimer of HSPB1 and characterize its NTR both when tethered to its ACD and when presented as short peptides that represent NTR sub-regions were surprisingly revealing.. Peptides representing sub-regions of the NTR display specific interactions with the ACD of HSPB1. Furthermore, PRE experiments revealed that the interactions have preferred orientations. In addition to the previously observed interaction of the IXI motif from the CTR with the β4/β8 groove, our results establish four distinct NTR/ACD interactions: 1) distal region with β4/β8 groove, 2) aromatic region with L3/4 and L5/6, 3) conserved region with dimer interface groove, and 4) boundary region with dimer interface groove (Fig. 8F). The fact that multiple HSPB1 regions can bind to a given groove or surface sets up a situation in which there are more potential binding elements than there are binding sites. This, in turn, creates a large combinatorial array of possible states within a dimer, and even more states within an oligomer. For example, each HSPB1 dimer contains two β4/β8 grooves and four interacting regions (two copies each of the distal region and the CTR IXI motif). Our data and modeling indicate that a given groove may be 1) empty, 2) bound by an inter-chain distal region, 3) bound by an intra-chain distal region, or 4) bound by a CTR IXI motif (usually from another dimer within an oligomer). A dimer may have zero, one, or two of its grooves filled, presumably with any combination of binders. The HDX results on HSPB1 oligomers indicate that a large proportion (roughly half) of the distal regions and the CTRs are bound, meaning the β4/β8 grooves must be predominantly occupied. Each HSPB1 dimer has a single dimer interface groove, but its potential interactions with two NTR regions creates a similarly complicated situation: a given dimer interface groove may be empty, bound by a single boundary region, a single conserved region, two boundary regions, one boundary plus one conserved, or two boundary regions plus a conserved region. Again, in the context of an oligomer, the combinatorial possibilities will be increased if the interactions can occur from neighboring dimer units.

While the NTR of sHSPs has long been known to be intrinsically disordered, it is clear from this work and recent crystal structures that further definition is required to accurately describe and define sHSP NTRs. Indeed, the only segment of the NTR that exhibits solvent-accessible, random coil-like behavior is the insertion sub-region. In recent years, it has come to be appreciated that the term intrinsic disorder can encompass a broad variety of behaviors beyond random coil. Disordered proteins or regions that adopt compact conformations and that transiently sample secondary structure have been described as molten globule-like, and disordered regions that interact with binding partners in a dynamic manner have been described as fuzzy^34, 35^. By definition, fuzzy protein-protein interactions cannot be described by a single conformational state^36^. However, given the high degree of orientational specificity of many NTR-ACD interactions, these interactions can be described neither as fuzzy in the canonical sense, nor as molten globule-like. The only region of the NTR which could be said to interact with the ACD in a fuzzy manner is the insertion region, as a spin label placed in this position causes non-specific PREs in many regions of the ACD, inconsistent with a single conformational state or orientation.

Notably, ordered interactions occur for several NTR sub-regions with the ACD with varying levels of affinity, and some interactions appear to be interdependent. The high degree of heterogeneity in HSPB1 dimers and oligomers is generated not by multiple random or fuzzy states but rather by the large number of possible combinations of several specific and orientationally-defined states. Based on observation of multiple slowly exchanging peaks by NMR for certain residues and bimodal HDXMS at long time points, the lifetimes for these interactions range from a minimum of tens of milliseconds to several minutes. For this reason, we propose the term “quasi-ordered” to describe the NTR of HSPB1, as it makes highly-specific long-lived (on the timescale of seconds) contacts while remaining dynamic and heterogeneous.

The structural information gleaned from the approaches presented here on HSPB1 add substantially to emergent structural insights on human sHSPs. First, knob-into-hole binding of CTR IXI motifs and β4/β8 grooves has been well established in numerous sHSPs. This type of interaction was recently reported for NTR IXI motifs as well^16, 20^. Recognition of non-canonical hydrophobic motifs by the β4/β8 groove has been shown previously for the client proteins amyloid-β and α-synuclein and in crystal structures of artificially-truncated sHSP constructs^37–41^, but never within a full-length sHSP. Our results show that a motif of alternating hydrophobic residues in the HSPB1 distal region competes effectively with the canonical IXI motif in the protein’s CTR, expanding the repertoire of potential binding partners.

The eight-residue conserved motif is the only stretch of identifiable sequence conservation in the NTR of human sHSPs. Recently, conserved motifs bound at the dimer interface groove have been observed in HSPB6 and HSPB2/3 structures. In the HSPB6 structure, the conserved sequence occupies the dimer interface groove and no β2 strand is present^20^. In the HSPB2/HSPB3 structure, the conserved motif of HSPB2 and one copy of a β2 strand from the same protomer occupy the groove. Our modeling shows that it is physically possible for two copies of β2 strand and a conserved motif to be bound simultaneously. The reciprocal effects observed in the HDX of disease-associated mutations in these two regions (G34R- and G84R-HSPB1) strongly suggest that disruption of one of the interactions destabilizes the other.

An interaction analogous to the one defined for the aromatic sub-region of HSPB1 has not been previously reported. Our results indicate that the interaction is favored in the absence of Ser15 phosphorylation, which disrupts the interaction. Only two other human sHSPs are enriched in aromatic residues in this region, namely HSPB4 and HSPB5. HSPB1, B4, and B5 are also the only human sHSPs known to form large oligomers, leading us to propose that the aromatic region/ACD interaction may be a driver of large oligomer formation. Notably, HSPB1 and HSPB5 are both phosphorylated in response to stress conditions on serine residues within their aromatic regions, yielding smaller oligomeric species, but the local mechanism by which this occurs had not been defined. Our results show clearly that serine phosphorylation disrupts the interaction in HSPB1, pointing to a shared structural mechanism by which stress-induced phosphorylation disrupts HSPB1 and HSPB5 oligomers, allowing them to form smaller, more dispersed, and more active species^25, 42^.

Intriguingly, PRE results place residue 83 in the same region where we observe aromatic peptide binding, indicating that the other two phosphorylation sites of HSPB1 (78 and 82) are in proximity to the aromatic region. Several sHSPs have phosphorylation sites in similar sub-regions, and it has been shown for HSPB1 that similar effects on quaternary structure and chaperone activity are obtained from mutation of any of these sites^29^. A recent crystal structure of an HSPB1 construct containing part of the boundary region and the ACD in complex with a peptide containing residues 76-88 phosphorylated at position 82 contained density in this same region. While this density was attributed to the peptide, the identity of the residues could not be resolved^37^. Our results confirm that the two phosphorylation sites could reside near this location. Thus, all three sites of stress-induced modification are near each other and close to a conserved ACD surface that is highly enriched in negatively-charged amino acids (i.e., L3/4 and L5/6), providing an environment that can easily be disrupted by additional negative charge. Both regions were less protected as seen by HDXMS in the phosphorylation-mimicking dimer compared to WT oligomers, consistent with coupled behavior. It has been demonstrated that singly- and doubly-phosphorylated HSPB1 species adopt intermediate-sized oligomers^29^, consistent with the arrangement acting as a rheostat that tunes the distribution of oligomeric states in response to cellular stress. Our results placing sequentially distant phosphorylation sites in spatial proximity demonstrates a common mechanism for the global effects observed for mutation of different phosphorylation sites. While there is clear interplay among the three phosphorylation sites, additional studies will be required to determine the nature of the interactions between the sites in the boundary region and the aromatic region. In particular, the role of boundary region phosphorylation is unclear, as we did not detect a difference in binding by the phosphorylated and non-phosphorylated forms of this peptide. Recent NMR work showed that an elongated ACD construct containing the boundary region with phosphomimetic mutations at regions 78 and 82 formed a transient β2 strand in solution^37^. However, without a direct comparison to a construct of the same length but without phosphomimetic mutations, the role of phosphorylation in this interaction remains unclear.

Our study also provides the first residue-level insights into the effects of known disease-associated mutations within the HSPB1 NTR. The three mutations investigated have previously been shown to alter the oligomeric distribution of HSPB1, in each case yielding larger oligomers^32^, but an understanding of the effects on a more detailed level are lacking. Remarkably, single mutations in the NTR have profound, widespread effects on dynamics, highlighting sHSP sensitivity to mutation and modification. We find that mutations at residues only five positions apart in the NTR have distinct, almost opposite effects (G34R and P39L) while two mutations that are 50 residues apart from each other (G34R and G84R) produce highly similar effects. In particular, G34R and G84R variants in the conserved and boundary regions respectively each exhibit a coupled increase in deuterium exchange in both the conserved and boundary regions. Furthermore, the mutant G34R conserved region peptide showed a lower affinity for the dimer interface groove. Altogether the results identify an interplay between two non-local regions of the NTR, in which the location of one region affects the other. Both regions can bind at the dimer interface groove, so another way to view the interdependence is that occupancy at a given interface groove by one sub-region favors occupancy by the other.

While the G34R and G84R substitutions are associated with the release of NTR sub-regions from their ACD interactions, P39L-HSPB1 shows markedly increased protection from exchange in the aromatic, conserved, and Trp-rich regions of the NTR. However, there is no large change for the boundary region, suggesting that the interdependence observed for the two glycine-to-arginine mutations is due to substitution of a bulky (and/or charged) amino acid in either the conserved or boundary regions *per se*. The increased helicity observed in the CD spectrum of P39L-HSPB1 oligomers is consistent with stabilization of helical structure in the Trp-rich region, likely a direct consequence of the helix-favoring proline-to-leucine substitution. Notably, no interactions were detected between the Trp-rich region and the ACD in either the WT- or mutant form, implying that the sub-region could be involved in NTR-NTR interactions. Our inability to make mutations in the Trp-rich sub-region that did not have an impact on oligomer size is further corroboration of the sub-region’s central role in driving HSPB1 oligomerization. Indeed, the decreased rate of subunit exchange from P39L-containing oligomers suggests that such NTR-NTR interactions play a rate-limiting role in the dissociation of subunits.

In sum, an approach using solution-state NMR, HDX/MS, and modeling has succeeded in defining the heretofore intractable NTR of HSPB1. Rather than behaving as a completely intrinsically disordered region, we find it to be quasi-ordered, with six sub-regions that display distinct properties and binding preferences. The results reveal that, contrary to expectation, the high degree of heterogeneity and polydispersity that is a defining feature of HSPB1 (and other human sHSPs) derives not from fuzzy disorder but rather from an array of combinatorial interactions that involve discrete NTR sub-regions and specific surfaces on the structured ACD. We expect other oligomeric sHSPs are similarly defined and that they can be parsed out using approaches similar to those described here. Finally, it is reasonable to think that there are other examples of quasi-order, with multiple interacting regions in a polypeptide chain, that can likewise be defined at a structural level by similar experimental approaches.

## EXPERIMENTAL PROCEDURES

### Protein expression and purification

Human HSPB1 (accession # P04792) had previously been cloned into pET23a and pET151d vectors (ampicillin resistant). The B1-ACD construct had previously been optimized to truncation of the full-length sequence from Gln80 to Ser176. Site-directed mutagenesis using the QuikChange protocol was used to introduce substitution mutations throughout the sequence.

Several protocols for protein expression were used to obtain different isotopically labeled samples. In almost all cases (unless otherwise specified) for full-length HSPB1, BL21(DE3) *E. coli* cells were used and a final concentration of 1.0 mM isopropyl β-D-1-thiogalactopyranoside (IPTG) was added to induce protein expression. For the B1-ACD construct, 0.5 mM IPTG was used.

For natural abundance (or non-isotopically labeled) protein, cells were grown in 0.5 L of lysogeny broth (LB) with 100 μg/mL ampicillin. Cells were grown at 37°C in a shaking incubator until OD_600_ ∼0.6. IPTG was then added to a final concentration of 1.0 mM and the temperature reduced to 22°C. Protein was expressed in a shaking incubator for ∼22 hours. Cells were harvested by centrifugation and resuspension in lysis buffer (50 mM Tris, pH 8.0, 100 mM NaCl, 1 mM ethylenediaminetetraacetic acid [EDTA]).

For ^15^N-labeled (no deuteration) protein, MOPS minimal media was used. Per 1 L culture, 1 g of ^15^NH_4_Cl was used for isotopic labeling. 4 g/L of glucose was used. Growth and expression steps were identical to those used for natural abundance protein.

For ^2^H^15^N^13^C-labeled (partial deuteration, ∼75%) protein, cells were grown in stages to acclimate the cells to deuterated minimal media. For all ^13^C-labeling, 3 g/L of ^13^C-glucose was used. One colony was grown in 3 mL of LB for ∼5 hours and then centrifuged to pellet the cells. These cells were resuspended and grown in 50 mL of H_2_O-based M9 minimal media to an OD_600_ ∼0.6. Cells were again pelleted, resuspended, and grown in 100 mL of D_2_O-based M9 minimal media to an OD_600_ ∼0.6. Cells were then transferred directly to a larger 500mL (total) D_2_O-based M9 minimal media and grown to an OD_600_ ∼0.6. After induction and reduction of temperature, protein was expressed for ∼48 hours.

For ^2^H^15^N^13^C-labeled (perdeuteration) protein, cells were grown in additional stages to acclimate the cells to deuterated minimal media. 3 g/L of ^2^H^13^C-glucose was used for the final culture. Stocks for deuterated M9 minimal media were also prepared in D_2_O. One colony was grown in 3 mL of LB for ∼5 hours and then centrifuged to pellet the cells. These cells were resuspended and grown in 50 mL of H_2_O-based M9 minimal media to an OD_600_ ∼0.6. Cells were again pelleted, resuspended, and grown in 100mL of D_2_O-based M9 minimal (non-deuterated glucose) media for 1-2 hours. Cells were again pelleted, resuspended, and grown in 200mL of D_2_O-based and ^2^H-glucose-based M9 minimal media to an OD_600_ ∼0.6. Cells were then transferred directly to a larger 1 L (total) D_2_O-based and ^2^H-glucose-based M9 minimal media. Cells were grown to an OD_600_ ∼0.6 and induced. After induction and reduction of temperature, protein was expressed for ∼48 hours.

Cells containing full-length HSPB1 were lysed by freeze-thaw and incubation with lysozyme and protease inhibitors in lysis buffer on ice for 20 minutes. Generally 0.5 L cultures of cells were lysed in one tube to maximize yield and purity. Deoxycholate was added to the lysed cells and placed on a shaking incubator at 37°C for 15 minutes. DNase, RNase, and magnesium chloride were then added and shaking incubation continued for 15 minutes. Cell lysate was centrifuged at high speed at 4°C. Ammonium sulfate was added to the supernatant to 40% saturation on a slow shaker at room temperature and allowed to equilibrate for 30 minutes. Ammonium sulfate precipitate was centrifuged at high speed at 20°C and the supernatant discarded. Pellets were used immediately or stored at −80°C for up to 1 week.

Ammonium sulfate pellets were resuspended in anion exchange buffer (AEX-20 mM Tris, 10 mM MgCl_2_, 30 mM NH_4_Cl, pH 7.6) at room temperature. Remaining solids were pelleted briefly at 20°C. Protein was desalted using a G25 column in AEX buffer at room temperature. Precipitated material was pelleted briefly at 20°C. Desalted protein was separated by an anion exchange DEAE column with a step gradient of AEX buffer with increasing sodium chloride at room temperature. Protein fractions were analyzed for purity by SDS-PAGE, pooled, and concentrated for SEC. Protein was separated on Superdex 200 or 75 columns in 50 mM sodium phosphate (NaPi), 100 mM NaCl, 0.5 mM EDTA, pH 7.5 buffer. Oligomeric proteins (WT, disease mutants) were separated on a Superdex 200 column, while smaller constructs (HSPB1_dimer_, NTR-ACD) were separated on a Superdex 75 column. SEC separated fractions were analyzed for purity by SDS-PAGE, pooled, and concentrated to the desired concentration. Concentration was determined by 280 nm absorbance (extinction coefficient of 40,450 M^-1^cm^-1^).

For cysteine-free proteins or proteins containing only the native cysteine at position 137, no reducing agent was added during purification. With the native cysteine present, the resulting protein was generally >95% oxidized at the dimer interface as seen by dimer formation by non-reducing SDS-PAGE. For proteins containing non-native cysteines (for fluorophore or spin labeling), the reducing agent dithiothreitol (DTT) was included at each stage of purification to avoid disulfide formation.

### Nuclear magnetic resonance spectroscopy

All NMR experiments were carried out on either 600 or 800 MHz Bruker spectrometers equipped with cryoprobes. All samples were prepared in 50 mM NaP_i_, 100 mM NaCl, 0.5 mM EDTA, pH 7.5 buffer. Spectra were collected at 30°C.

Several TROSY-based triple resonance experiments were implemented to assign peaks in the NTR-ACD spectrum to particular residues-HNCO, HN(CA)CO, HNCA, HN(CO)CA, HNCACB, HNCB, and HNCOCANNH (“NNH”) experiments. Non-uniform sampling (NUS) at a sampling rate of 25% was used for longer experiments. For NUS datasets, an iterative soft threshold algorithm was used to reconstruct full spectra^43^. A maximum protein concentration of 600 μM was used as inter-dimer interactions were evident at higher concentrations (many peaks broadened). Both ∼75% deuterated and perdeuterated ^2^H^15^N^13^C-labeled forms of NTR-ACD (expression in previous section) were used for almost all triple resonance experiments, with modest improvements observed for higher deuteration levels. For the HNCB experiment, only perdeuterated protein was used.

To measure intensities and positions of peaks in 2D spectra, NMRViewJ was used^44^. The following equation was used to calculate chemical shift perturbations (CSPs) between peaks in two spectra:

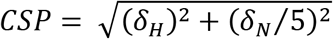

Spin-label constructs were made with a NTR-ACD/C137S background and cysteines introduced at positions throughout the NTR, one at a time. Samples were prepared in 50 mM NaPi, 100 mM NaCl, 0.5 mM EDTA, pH 7.5 buffer and 10 mM DTT to fully reduce all cysteines. Reducing agent was then removed from protein samples using a desalting column. Immediately after removing reducing agent, 5-fold molar excess (1-oxyl-2,2,5,5-tetramethylpyrroline-3-methyl)-methanethiosulfonate (MTSL) spin label (in DMSO) was added to protein samples and allowed to incubate overnight at 4°C or at room temperature for 2 hours. Excess MTSL was then removed from the labeled protein using a desalting column. The resulting NMR samples contained 300-400 μM protein. 2D ^1^H-^15^N HSQC-TROSY spectra were collected for each spin-labeled mutant protein. The unpaired spin label was then quenched in each sample by addition of ascorbate (5 mM final concentration). Identical spectra were collected for each quenched sample for intensity comparison between quenched and unquenched spectra.

The distal, aromatic, conserved, and boundary region peptides were purchased from Genscript. The N-terminal residue of all but the distal peptide was formylated, and the C-terminal residue of all peptides was amidated. The Trp-rich peptides were purchased from LifeTein and were acetylated on the N-terminus and amidated on the C-terminus. The distal, conserved, and boundary peptides were dissolved in NMR buffer prior to use. The aromatic and Trp-rich peptides were dissolved in DMSO to a concentration of 100 or 50 mM, then diluted in buffer to a concentration of 1 mM. All peptides were stored at −80°C. A spectrum of ^15^N B1-ACD in the presence of 1% DMSO was collected and used as a reference for the aromatic and Trp-rich peptide experiments, although it was highly similar to the ^15^N B1-ACD spectrum in the absence of DMSO.

### Hydrogen-deuterium exchange mass spectrometry

200 μM protein samples were equilibrated at room temperature (22°C) in 50 mM NaPi, 100 mM NaCl, 0.5 mM EDTA at pH 7.5 for several hours. Samples were diluted 10-fold into deuterated buffer (prepared identically but with D_2_O) for a final concentration of 20 μM and incubated at room temperature for various periods of time to allow for hydrogen-deuterium exchange. At the desired time point, the deuteration reaction was quenched by adding an equal volume of quench buffer (0.6% formic acid) on ice for a final pH of 2.5. Quenched samples were immediately flash frozen in liquid nitrogen and stored at −80°C. Undeuterated samples were prepared in a similar fashion but replaced addition of D_2_O buffer with protonated buffer. Fully deuterated samples were made by first denaturing the protein (3M guanidine HCl and high heat for at least 30 minutes), making the same dilution into D_2_O buffer, incubating for several hours, and quenching the same as all other samples.

Samples were stored in liquid nitrogen until 5 minutes prior to injection to maintain consistent levels of deuterium loss (back-exchange). Initially, the sample was passed over a custom packed pepsin column (1 x 50mm) at 200uL/min in 0.1% formic acid at 1°C for digestion of the protein into peptides. Digested peptides were then captured onto a trapping column (Waters vanguard BEH C18 2.1 x 5 mm 1.7 µm 130Å) and resolved over a C18 reverse-phase column (Waters BEH 1 x 100 mm 1.7 µm 130Å) using a linear gradient of 3 to 40%B over 10 minutes (A: 0.1% formic acid, 0.025% trifluoroacetic acid, 2% acetonitrile; B) 0.1% formic acid in acetonitrile). The LC system was coupled to a Waters SYNAPT G2 QTOF. The source and desolvation temperatures were 70 and 130°C, respectively. The StepWave ion guide settings were set to minimize non-uniform deuterium loss during desolvation^45^. The pepsin, trap, and resolving columns were washed extensively to reduce sample carryover^46, 47^. The resulting levels of carryover were below 5% for each peptide analyzed based on blank runs. Peptic peptides of WT and mutant proteins were analyzed by tandem MS (MS^E^) analyzed by ProteinLynx Global SERVER.

Most peptides could be directly compared among all mutants. For peptides containing a mutation the comparisons are only qualitative as the change in exchange could be an effect of both intrinsic exchange and local structure. MassLynx software was used to align spectra of various time points at the appropriate retention times, and HX-Express v2^48^ was used to analyze deuterium incorporation and perform bimodal analysis. The deuteration level at each time point was calculated relative to the deuteration levels of the undeuterated and fully deuterated spectra for each peptide. In cases where the fit was very poor due to very broad isotope distributions or clear bimodals, an alternative fitting was used to obtain a bimodal distribution, representing two distinct deuteration states of the peptide at a given time point. For bimodal peptides shown in Figures 7 and 9, their behavior was confirmed across several charge states and in most cases several overlapping peptides. Three biological replicates of WT oligomers and HSPB1_dimer_ were examined and showed qualitatively similar patterns in HDX (Fig. 7 Supp2), with one of these replicates containing the native C137 and including reducing agent to confirm similar behavior to C137S constructs. Disease mutants were examined with single replicates due to limitation of instrument time.

### Modeling

A starting homology model was produced by aligning a crystal structure of the HSPB1 ACD (PDB 4MJH) with a crystal structure of an HSPB2/HSPB3 heterotetramer (PDB 6F2R) and creating a new PDB file with structural elements from both. It contained the HSPB1 ACD, including both β2 strands, a copy of the HSPB2 motif ^157^VNEVYISLL^164^ bound in each β4/β8 groove, and a copy of the HSPB2 conserved motif ^23^RLGEQRFG^30^ bound in the dimer interface groove. Residues from the HSPB2/HSPB3 crystal structure were mutated to the appropriate sequence for HSPB1 using the PyMOL Molecular Graphics System (Version 2.0, Schrödinger LLC). PyRosetta^49^ was used to eliminate clashes and model in missing segments. The function FastRelax^50^ was used to relax the starting homology model and eliminate clashes between chains. Fragment insertion using the BlueprintBDR^51^ mover was then used to add the initial residues of the distal region and a segment of the 2/3 loop that was not resolved in the crystal structure. Two distinct connectivities involving the aromatic region were modeled. One structure was generated in which the aromatic region connected the distal region bound in the opposite orientation of the conserved region, such that it wrapped around the edge of the ACD and contacted loops 3/4, 5/6 and 8/9. Another was generated in which it connected the conserved region to the other distal region and crossed over strands β8, β9, and β3. An initial model for each of these two structures was generated using the BlueprintBDR mover, and then subject to 100 rounds of Generalized Kinematic Closure^52^ to generate two sets of 100 structures that sample the conformational flexibility of the aromatic region. Generalized Kinematic Closure was used to model this longer disordered segment because, unlike the BlueprintBDR mover, it does not explicitly bias structures toward a smaller radius of gyration or use stretches of PDB-derived torsion angles. The missing loops connecting the beginning of the NTR to the β2 strands were then modeled in using the BlueprintBDR mover. Again, there were two possible connectivities for each structure: the conserved motif could be connected to either β2 strand, while the distal region not connected to the conserved motif could be connected to the other β2 strand. For each model, both connectivities were sampled, producing four sets with unique topologies that each contained 100 structures. For each step in which additional residues were introduced in the model, FastRelax was used to relax the new segment and the residues adjacent to it. Finally, FastRelax was used to relax all atoms in the final structures.

### Circular dichroism spectroscopy

Samples were prepared in 25 mM NaPi, 50 mM NaCl, and 0.25 mM EDTA buffer at pH 7.5. Samples were incubated at 20 μM at room temperature for several hours prior to measurement. All measurements were made on a Jasco J-1500 CD spectrometer with Peltier temperature control at 20°C, with 1 nm bandwidth, and averaged over three scans. All data were normalized to units of mean residue ellipticity (MRE). Data were collected at 0.1 nm intervals and then smoothed linearly across 1 nm. All data presented have a high-tension voltage below the recommended cutoff for the detector (800 V).

### Fluorescence-based subunit exchange

WT and disease mutant constructs were generated with a cysteine introduced at position 174, analogous to similar fluorescence studies in HSPB5^53^, and the native cysteine at position 137 mutated to serine. Protein was buffer exchanged into 50 mM NaPi, 100 mM NaCl, 0.5 mM EDTA, pH 7.5 buffer with 2 mM tris-(2-carboxyethyl)-phosphine (TCEP). Protein was incubated with 3X molar excess of Alexa Fluor 488 maleimide and incubated at 37°C (to facilitate subunit exchange) for at least one hour. Protein was separated from free dye using gravity desalting columns. Fractions collected from desalting columns were analyzed for separation by comparing absorbances at 495 nm (fluorophore extinction coefficient of 73,000 M^-1^cm^-1^) and 280 nm, with an assumed A_280_/A_495_ ratio of 0.11. The resulting pool of fractions was diluted to desired protein concentrations and fluorophore labeling percentages.

Samples were incubated at 37°C at 20 μM and labeling percentage for at least one hour prior to measurement. For the experiments presented here, samples with 30% fluorophore labeling were mixed 1:2 with unlabeled protein. The resulting dequenching from homo-FRET among fluorophores in oligomers was measured as a function of time. Measurements were collected on a Horiba Fluorolog-3 with double excitation and emission monochromators and Peltier temperature control at 37°C. Excitation and emission wavelengths of 518 nm and 498 nm were used, respectively.

The resulting kinetic data was fit to the following exponential equation using the R software package to obtain subunit exchange rates:

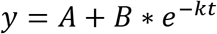

## ACKNOWLEDGEMENTS

We thank Ponni Rajagopal for extensive training and discussions of NMR experiments. We thank Jianming Kang for assistance with HPLC. Finally, we thank members of the Klevit lab for useful discussions and suggestions. This work was supported by NIH grant 2R01 EY017370 to R.E.K, R01GM127579 to M.G., NIH T32 GM008268 and Hurd Fellowship in Biophysics from the UW School of Medicine to A.F.C. and H.E.R.B., and the Hope Barnes Graduate Fellowship from the UW School of Pharmacy to H.E.R.B.

## SUPPLEMENTAL INFORMATION

**Figure 1 Supplement.**
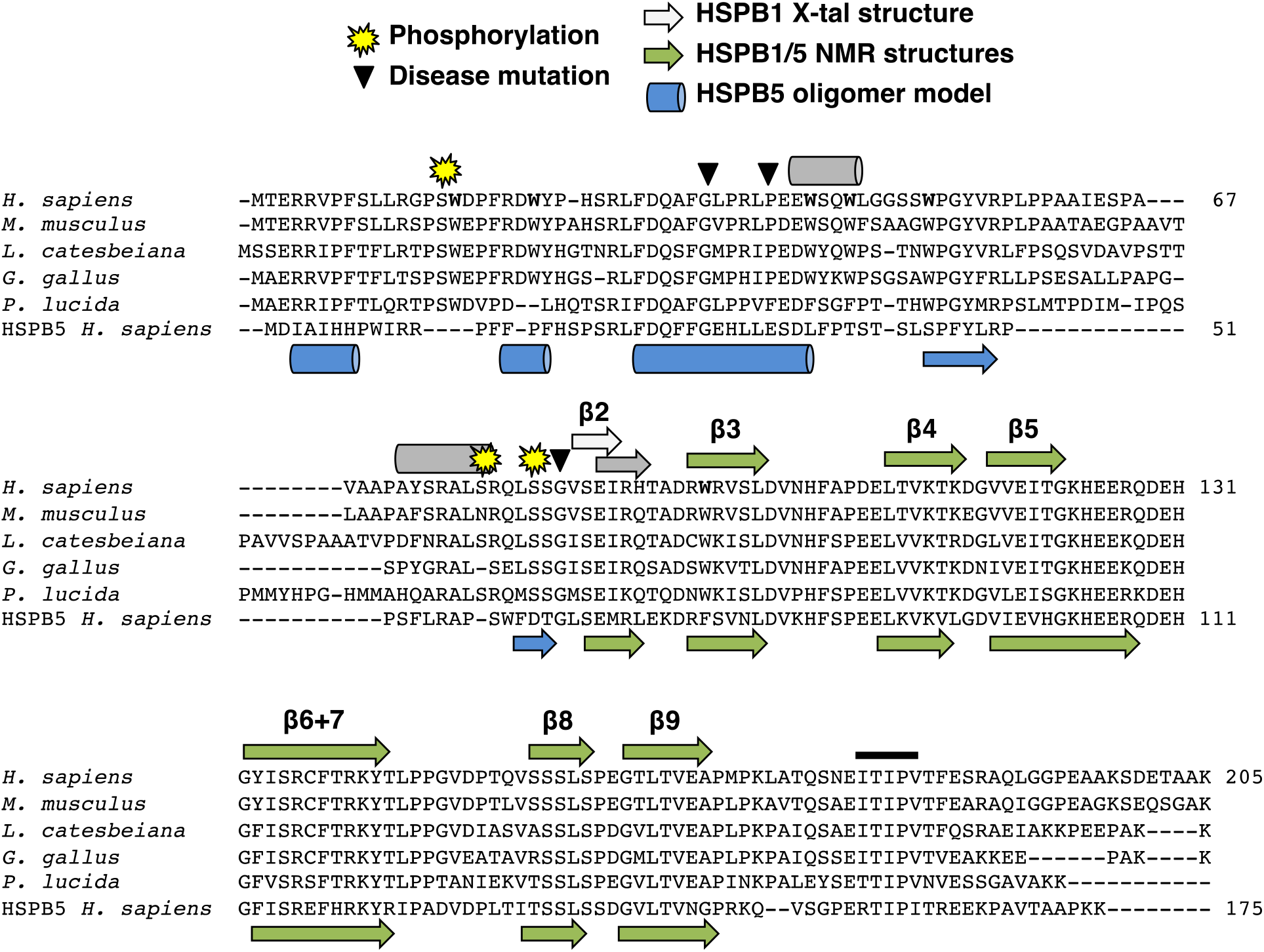
Sequence alignment of HSPB1 orthologs, residues of interest, and known and predicted secondary structure. HSPB1 sequences from human, mouse, frog, chicken, and fish show considerable conservation throughout most of the NTR, in contrast to other sHSP types as seen in the human HSPB5 sequence at the bottom. Yellow stars indicate conserved phosphorylation sites (S15, S78, and S82), and triangles indicate disease-associated point mutations in the NTR examined in this study (G34R, P39L, and G84R). Gray helices and strands on top of the sequence are based on PSIPRED secondary structure prediction^54^. The light gray β2 strand is based on the HSPB1 ACD-only crystal structure (PDB ID: 4MJH). The blue helices and strands below the sequence are based on secondary structure in the HSPB5 oligomeric model (PDB ID: 3J07). Green strands (below and above sequence) are based on HSPB1 and HSPB5 NMR ACD-only structures (PDB IDs: 2N3J and 2N0K). The black bar above the CTR indicates the IXI motif in HSPB1, which is actually a triple motif (ITIPV).

**Figure 2 Supplement 1.**
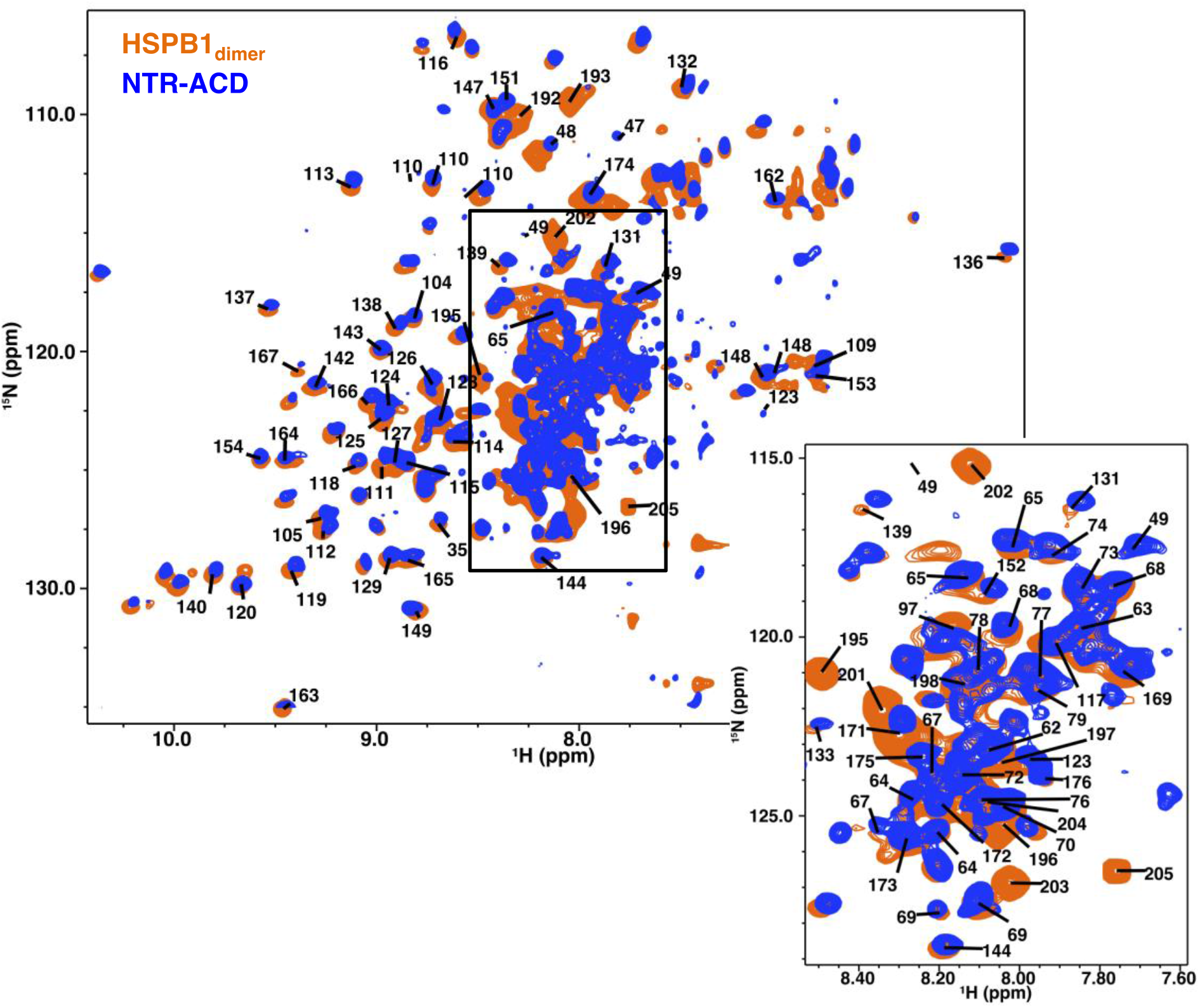
NMR spectra comparison of HSPB1_dimer_ and NTR-ACD. ^1^H-^15^N HSQC-TROSY spectra of the two constructs, showing considerable overlap; strong peaks in the center of the spectrum correspond to disordered regions of the protein (NTR and CTR). Center peaks are omitted for clarity and can be seen in the zoomed panel, which is at a higher noise level to better separate strong peaks.

**Figure 2 Supplement 2.**
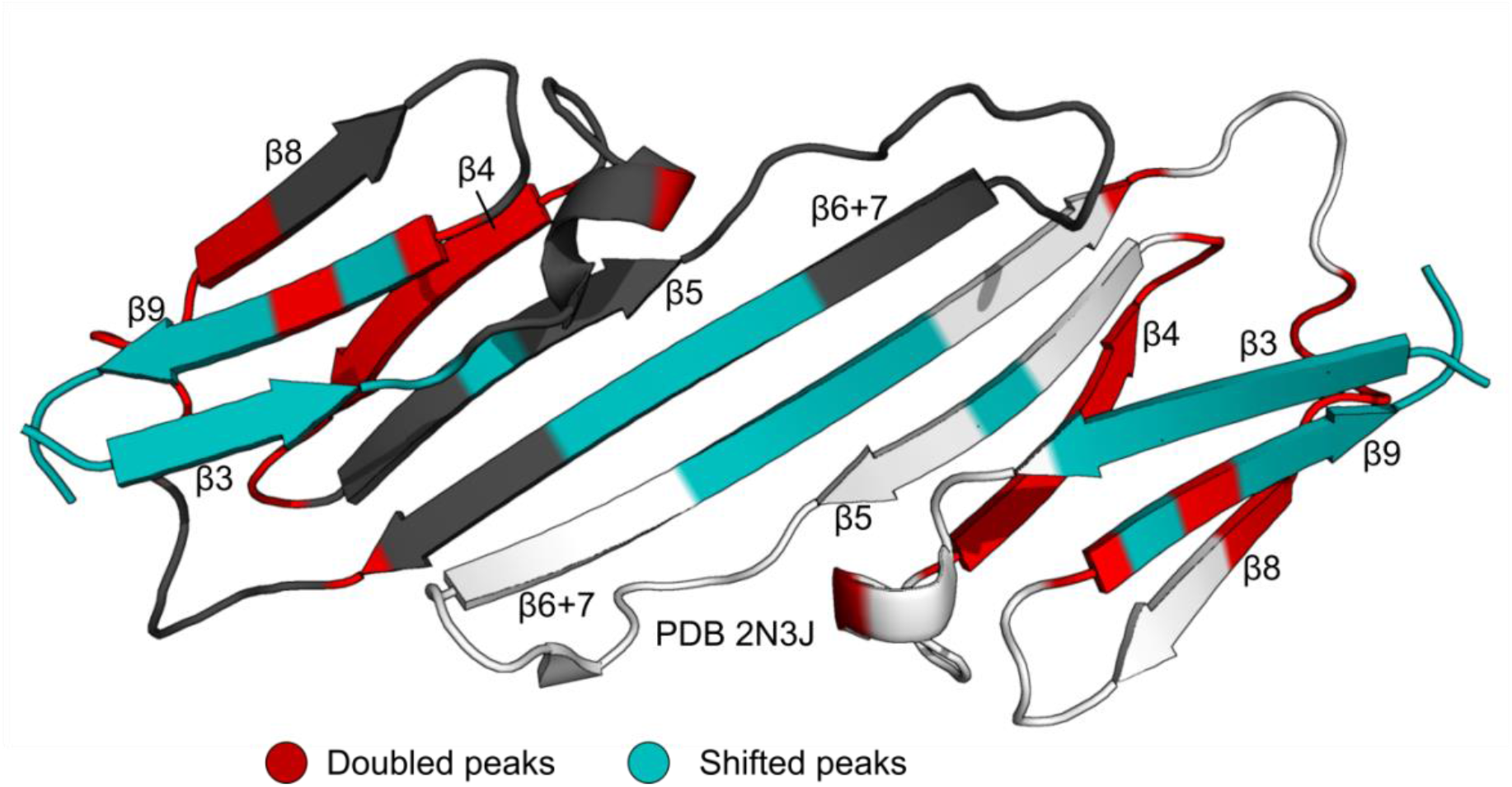
Perturbed residues in the mixed ^15^N-B1-ACD/NTR-ACD dimer spectrum. Residues for which a distinct, new peak arises in the same position as the NTR-ACD spectrum are highlighted in red. These mainly cluster around the β4-β8 groove. Residues for which the peak is perturbed, but there is not a clear, new peak are highlighted in teal. These occur in and around the dimer interface.

**Figure 4 Supplement.**
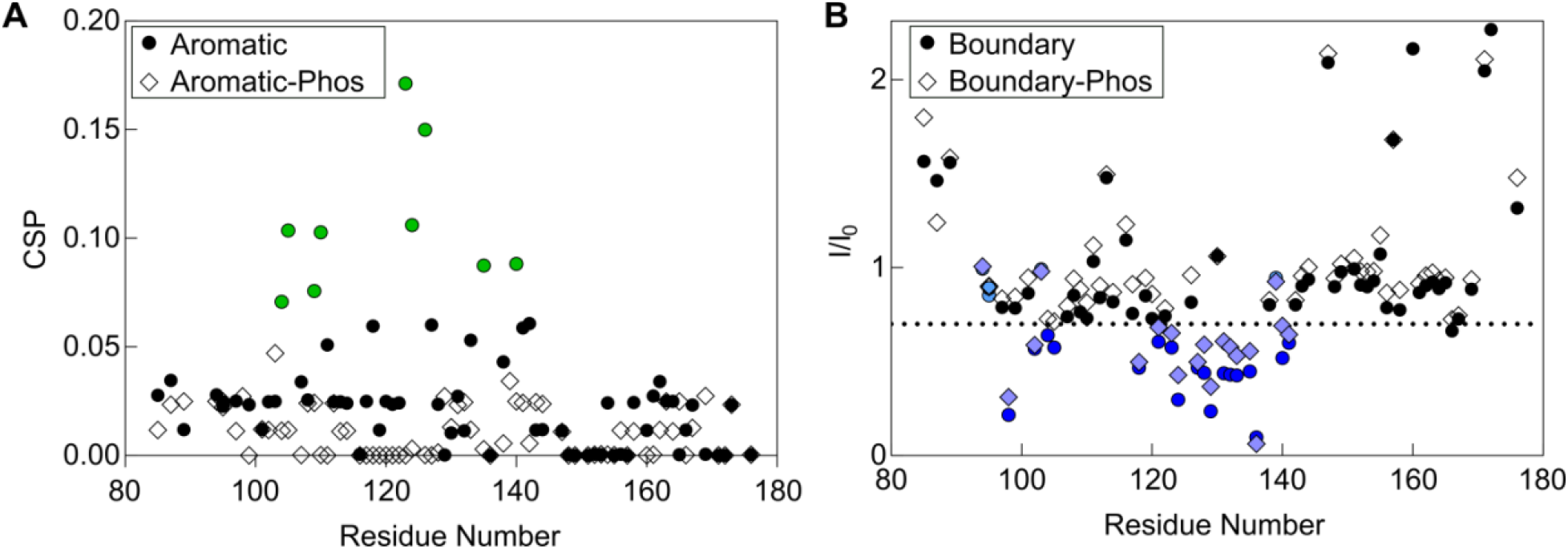
Effect of phosphorylation on peptide binding. (A) CSPs in the aromatic peptide binding experiment with ^15^N B1-ACD. A phosphorylated form of the aromatic peptide with phophoserine at position 15 does not cause strong CSPs (diamonds), indicating that phosphorylation disrupts binding of this peptide to the ACD. (B) NMR intensity loss in boundary region peptide binding experiment with ACD. A peptide with phosphoserine at positions 78 and 82 caused similar perturbations to the non-phosphorylated peptide, indicating that phosphorylation does not have a direct effect on this interaction. Colored points lost >30% intensity, had CSPs over 0.5, or both.

**Figure 7 Supplement 1.**
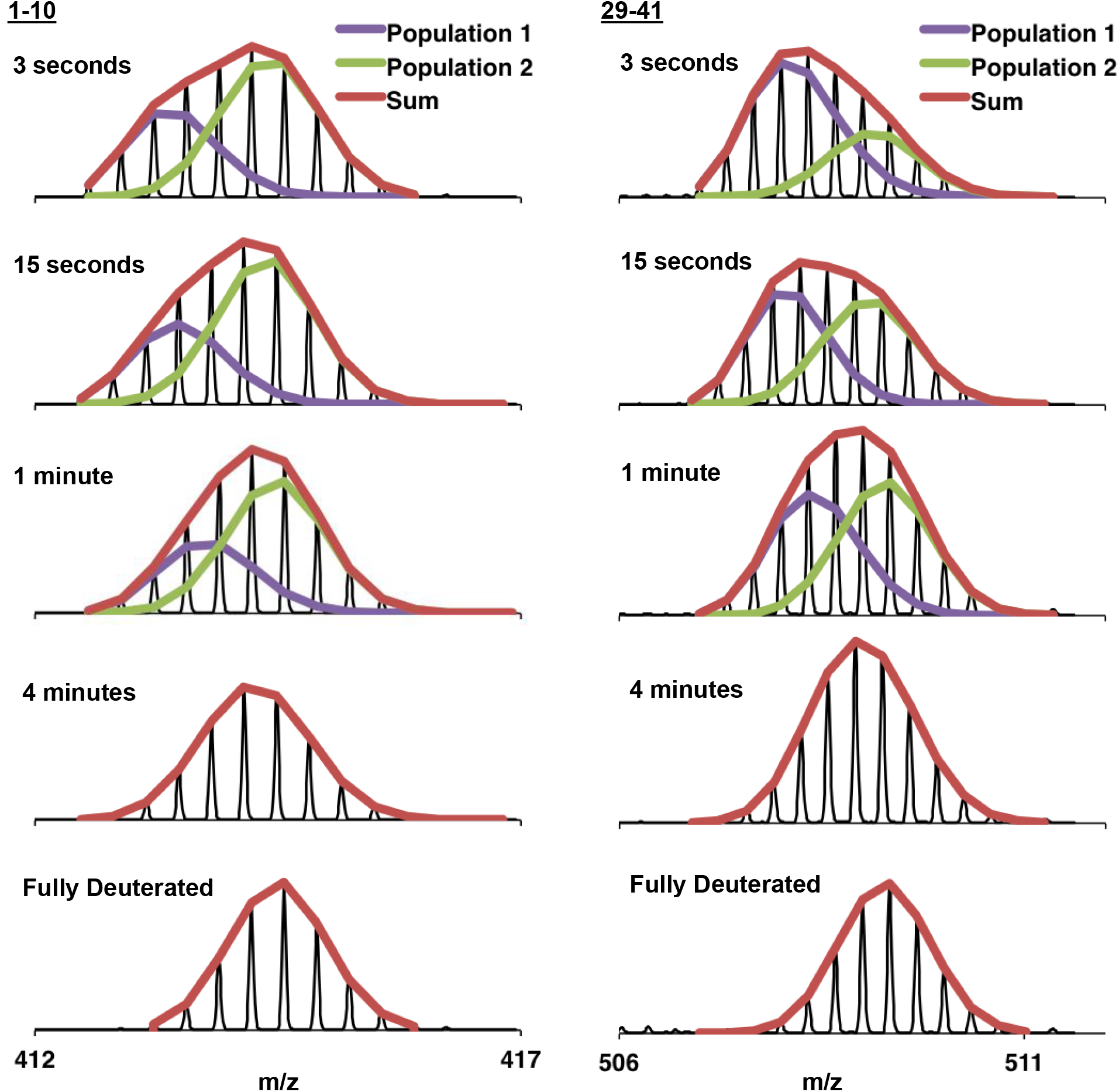
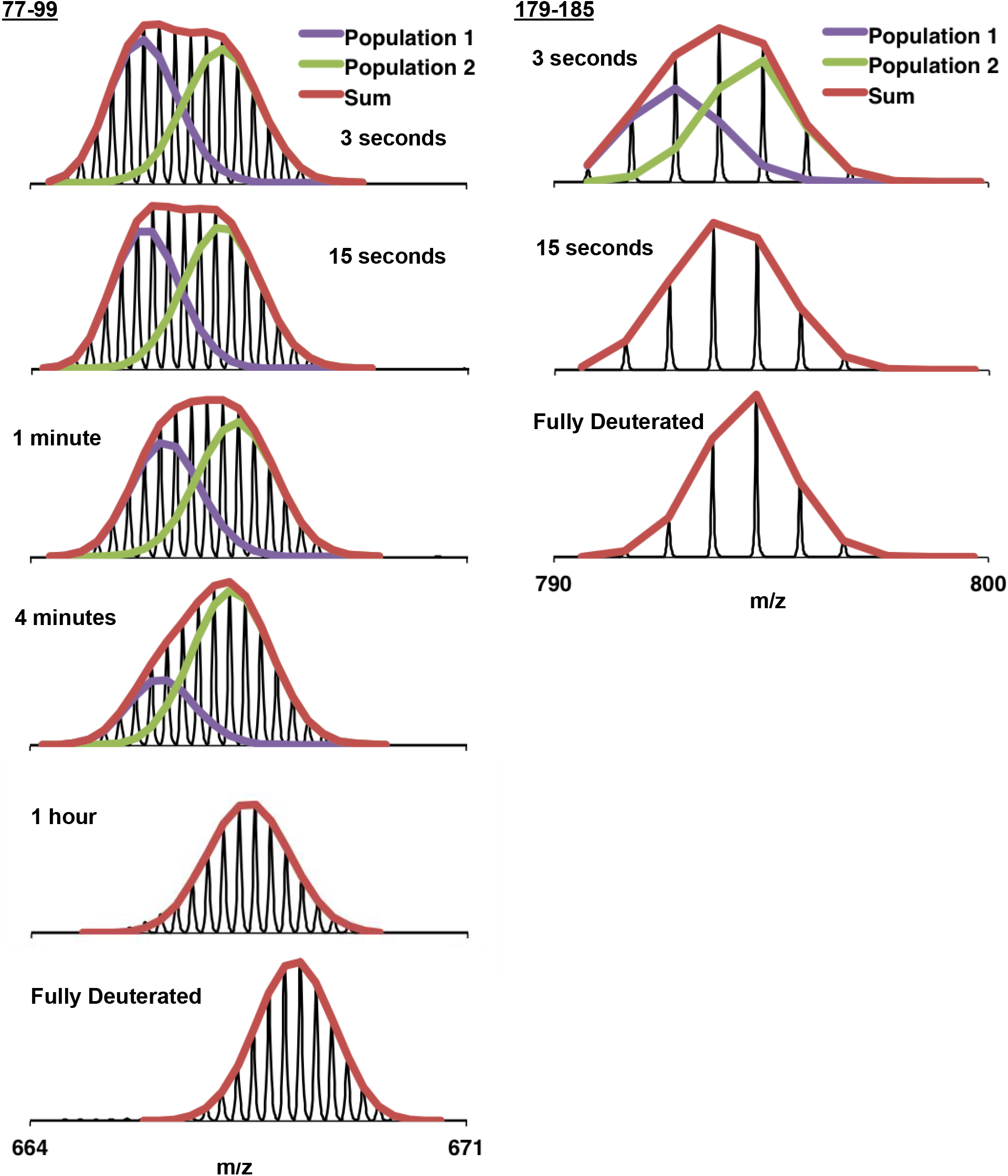
Example bimodal spectra for peptides 1-10, 29-41, 77-99, and 179-185 of C137S oligomers. Multiple populations (bimodals) are observed in earlier timepoints for several regions of the protein. HX-Express^48^ was used to deconvolute the two populations into two binomial fits. The sum of the binomial fits are shown in red.

**Figure 7 Supplement 2.**
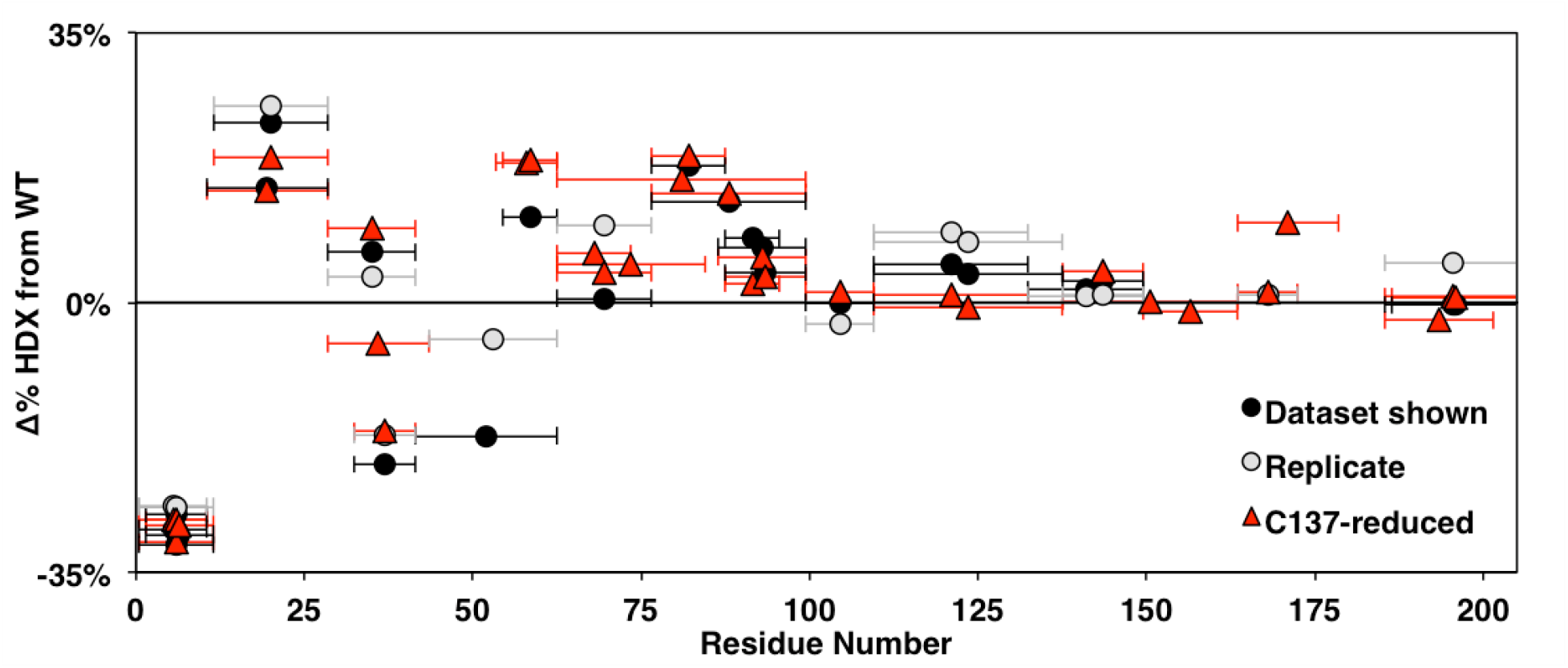
Comparison of changes in HDX between WT and HSPB1_dimer_ across biological replicates at 3 seconds. Identical constructs expressed, purified, and examined separately are shown in black and gray. The native C137-containing constructs with reducing agent are shown in red. Points in the upper half indicate decreased protection in HSPB1_dimer_ and points in the lower half indicate increased protection relative to WT oligomers. Qualitatively similar patterns are observed throughout the NTR.

**Figure 8 Supplement.**
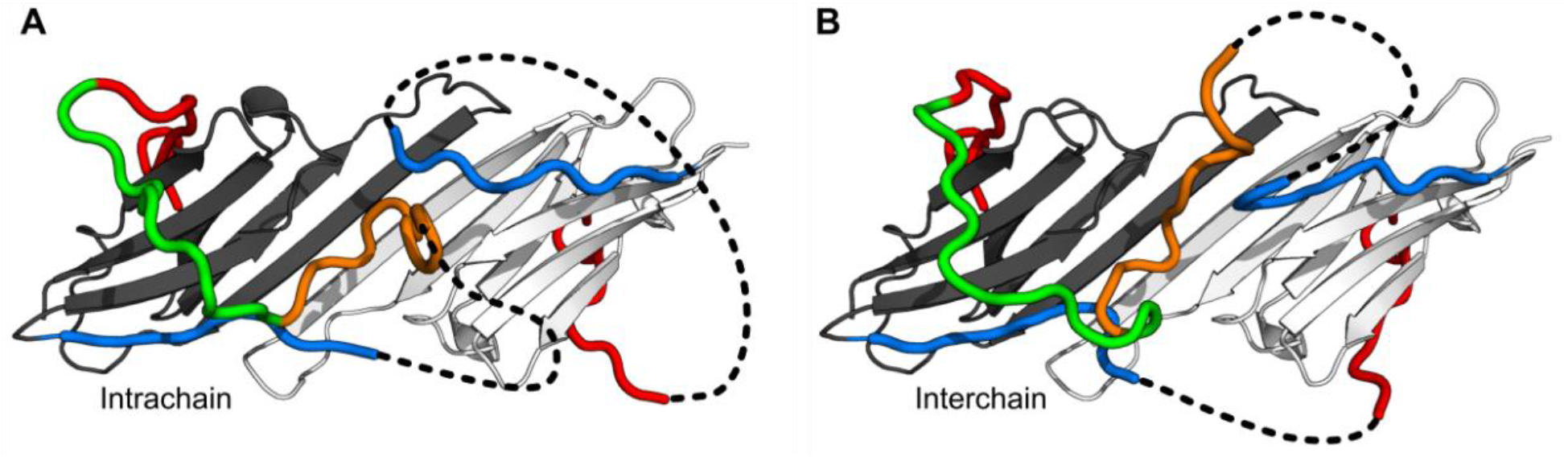
Models in which the conserved motif is connected to the distal motif oriented in the opposite direction. While we believe this connectivity is less favored than the models shown in the main figure, we were able to generate physically realistic models. Depending on which copy of β2 the conserved motif is connected to, contacts between the distal and aromatic regions and the ACD can occur intrachain (A) or interchain (B).

**Figure 9 supplement.**
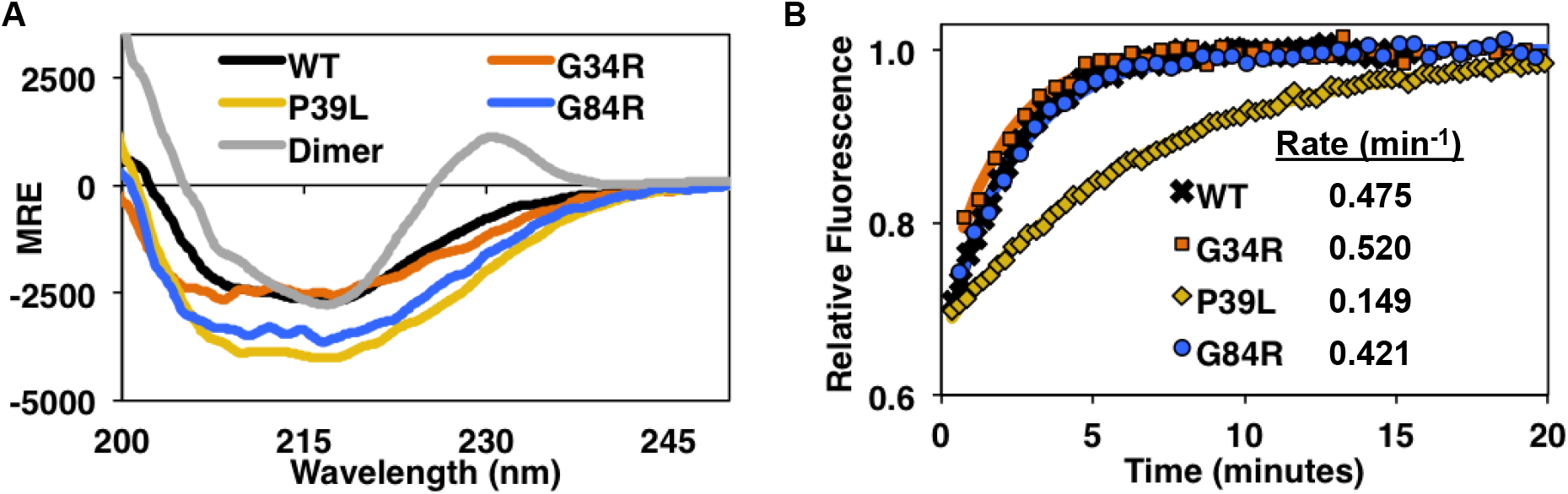
Circular dichroism spectra and subunit exchange kinetics of disease mutants. (A) Far-UV CD spectra shows overall similar secondary structure for WT, G34R, P39L, and G84R mutants, but P39L shows the strongest increase in ellipticity. The HSPB1_dimer_ has a unique positive peak at 230 nm as previously reported (Baughman). (B) Fluorescence-based subunit exchange of mutant oligomers at 37°C. AlexaFluor-488 labeled oligomers (fluorescence quenched) are mixed with unlabeled oligomers. As subunits exchange among oligomers, fluorescence increases as quenching is released. Only P39L has a dramatically decrease in the subunit exchange rate.

**Figure 7 Supplemental Table 1:**
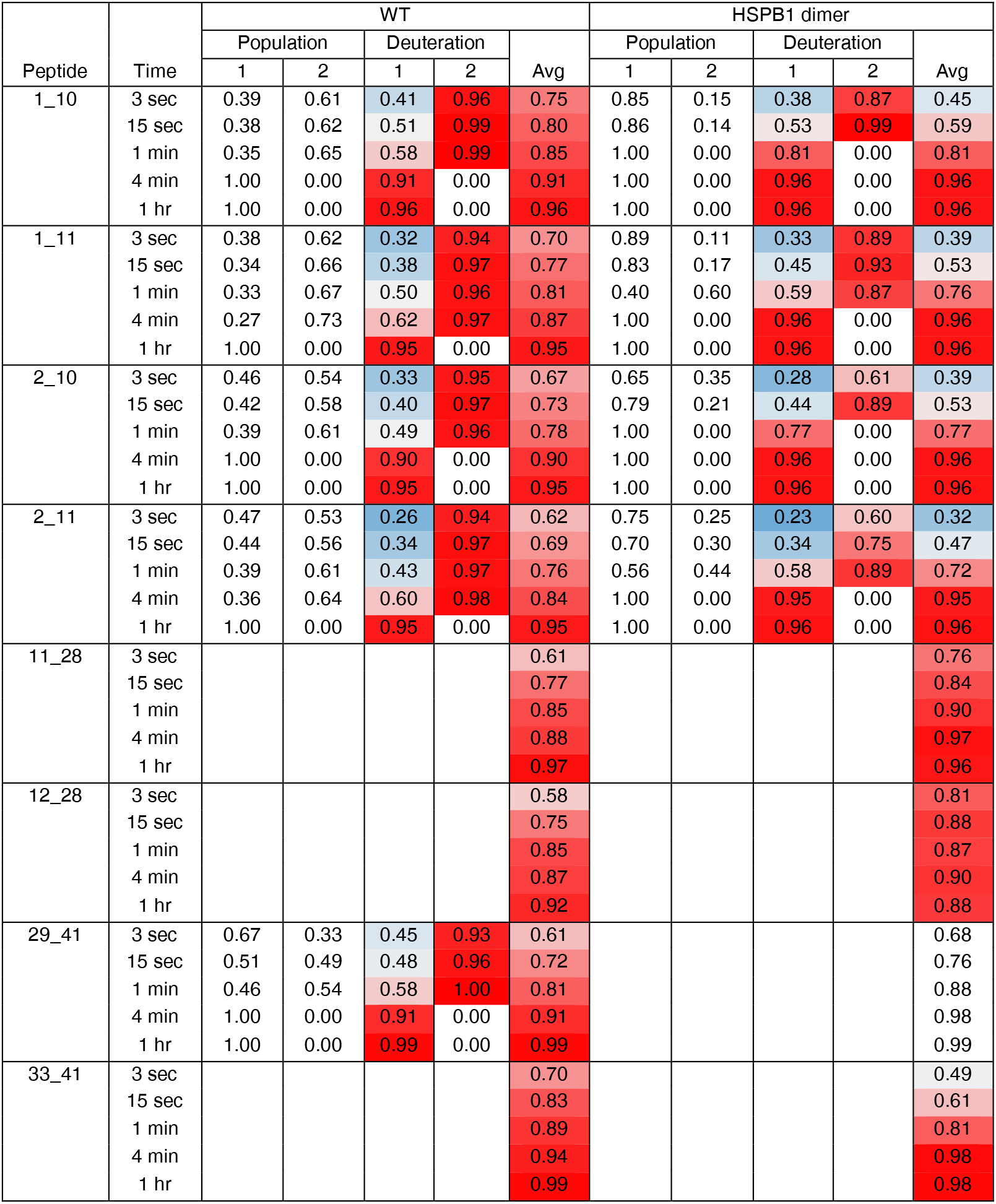
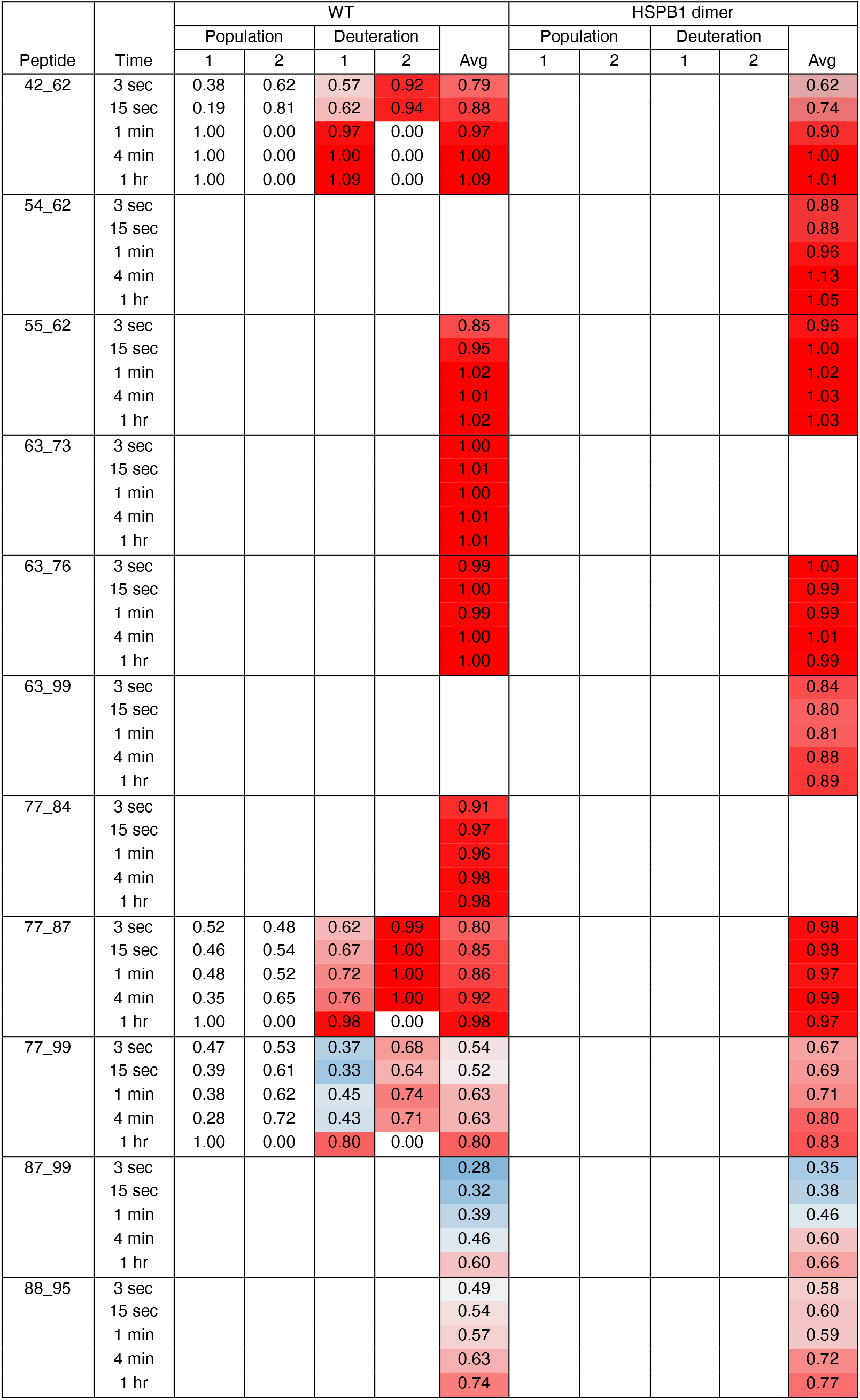
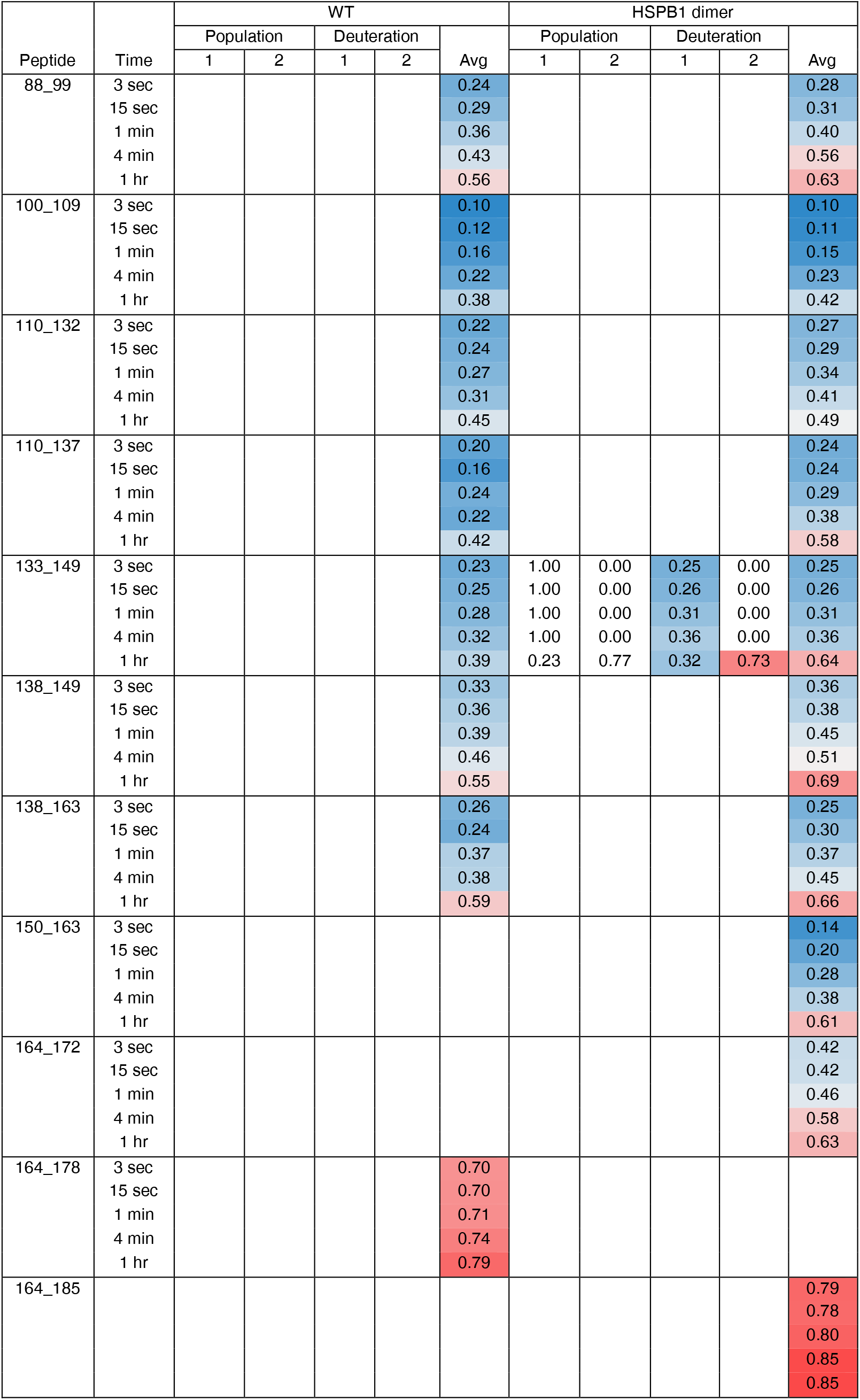
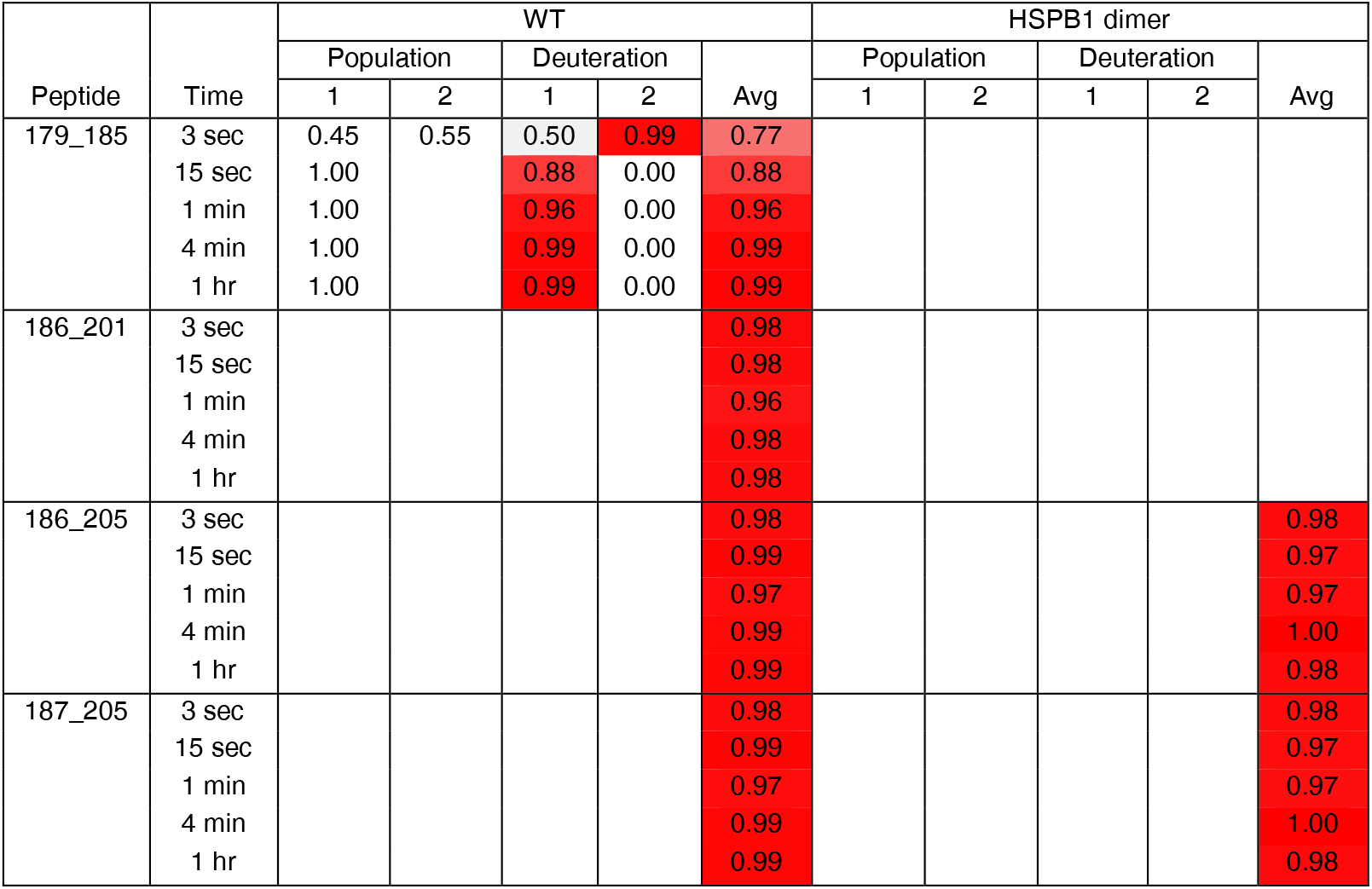
HDXMS profiles for WT and HSPB1_dimer_ forms of HSPB1. The deconvoluted fractions of each population from bimodal distributions are shown in the first two columns for each mutant. The fractional deuterium uptake for each of these populations is shown in the next two columns, followed by a weighted average in the fifth column. All spectra were binomial fit using HX-Express^48^.

**Figure 7 Supplemental Table 2.** Statistics for HDXMS experiments on all mutants. Statistics listed are for C137S forms of the following mutants of HSPB1. Sequence coverage takes into account the lack of observed deuterium uptake in the first two residues of each peptide. Back-exchange is calculated as the % deuterium uptake of the fully deuterated sample divided by the theoretical maximum deuteration for each peptide (89%), excluding prolines and the first two residues.

| Dataset/construct | WT | HSPB1<sub>dimer</sub> | G34R | P39L | G84R |
| --- | --- | --- | --- | --- | --- |
| Rxn conditions | Room temp. (25°C), pH 7.5, diluted 10X for final 89% D<sub>2</sub>O |  |  |  |  |
| Time points | 3 sec, 15 sec, 1 min, 4 min, 1 hour, fully deuterated |  |  |  |  |
| # of peptides | 29 | 29 | 38 | 36 | 34 |
| Sequence coverage | 90% | 93% | 93% | 94% | 93% |
| Redundancy | 1.8 | 1.9 | 2.5 | 2.4 | 2.2 |
| % Back-exchange (mean/IQR) | 24.6/14.0 | 24.5/12.9 | 20.4/13.8 | 20.3/13.6 | 19.2/14.7 |
| Replicates | Single, representative of 3 biological replicates |  | Single |  |  |
| Significance threshold | Qualitative assessment of differences > 5% exchange from WT |  |  |  |  |

**Figure 9 Supplemental Table:** HDXMS profiles for G34R, P39L, and G84R forms of HSPB1. The deconvoluted fractions of each population from bimodal distributions are shown in the first two columns for each mutant. The fractional deuterium uptake for each of these populations is shown in the next two columns, followed by a weighted average in the fifth column. All spectra were binomial fit using HX-Express^48^.

| Peptide | Time | G34R |  |  |  |  | P39L |  |  |  |  | G8R |  |  |  |  |
| --- | --- | --- | --- | --- | --- | --- | --- | --- | --- | --- | --- | --- | --- | --- | --- | --- |
|  |  | Population |  | Deuteration |  | Avg | Population |  | Deuteration |  | Avg | Population |  | Deuteration |  | Avg |
|  |  | 1 | 2 | 1 | 2 |  | 1 | 2 | 1 | 2 |  | 1 | 2 | 1 | 2 |  |
| 1_10 | 3 sec | 0.37 | 0.63 | 0.41 | 0.97 | 0.76 | 0.52 | 0.48 | 0.43 | 0.96 | 0.69 | 0.43 | 0.57 | 0.40 | 0.93 | 0.70 |
|  | 15 sec | 0.34 | 0.66 | 0.54 | 0.98 | 0.84 | 0.45 | 0.55 | 0.53 | 0.97 | 0.77 | 0.32 | 0.68 | 0.51 | 0.93 | 0.80 |
|  | 1 min | 0.27 | 0.73 | 0.62 | 0.98 | 0.88 | 0.46 | 0.54 | 0.65 | 0.97 | 0.82 | 0.16 | 0.84 | 0.56 | 0.90 | 0.84 |
|  | 4 min | 1.00 | 0.00 | 0.95 | 0.00 | 0.95 | 1.00 | 0.00 | 0.89 | 0.00 | 0.89 | 1.00 | 0.00 | 0.91 | 0.00 | 0.91 |
|  | 1 hr | 1.00 | 0.00 | 0.96 | 0.00 | 0.96 | 1.00 | 0.00 | 0.94 | 0.00 | 0.94 | 1.00 | 0.00 | 0.95 | 0.00 | 0.95 |
| 1_11 | 3 sec | 0.34 | 0.66 | 0.32 | 0.93 | 0.73 | 0.44 | 0.56 | 0.31 | 0.89 | 0.64 | 0.38 | 0.62 | 0.28 | 0.88 | 0.65 |
|  | 15 sec | 0.25 | 0.75 | 0.39 | 0.96 | 0.82 | 0.36 | 0.64 | 0.39 | 0.94 | 0.74 | 0.30 | 0.70 | 0.38 | 0.94 | 0.77 |
|  | 1 min | 0.23 | 0.77 | 0.51 | 0.97 | 0.87 | 0.29 | 0.71 | 0.46 | 0.94 | 0.80 | 0.29 | 0.71 | 0.50 | 0.95 | 0.82 |
|  | 4 min | 1.00 | 0.00 | 0.96 | 0.00 | 0.96 | 0.22 | 0.78 | 0.53 | 0.95 | 0.86 | 1.00 | 0.00 | 0.92 | 0.00 | 0.92 |
|  | 1 hr | 1.00 | 0.00 | 0.97 | 0.00 | 0.97 | 1.00 | 0.00 | 0.94 | 0.00 | 0.94 | 1.00 | 0.00 | 0.95 | 0.00 | 0.95 |
| 2_10 | 3 sec | 0.40 | 0.60 | 0.31 | 0.96 | 0.70 | 0.50 | 0.50 | 0.31 | 0.93 | 0.62 | 0.43 | 0.57 | 0.27 | 0.92 | 0.64 |
|  | 15 sec | 0.37 | 0.63 | 0.44 | 0.98 | 0.78 | 0.42 | 0.58 | 0.40 | 0.95 | 0.72 | 0.34 | 0.66 | 0.36 | 0.94 | 0.74 |
|  | 1 min | 0.31 | 0.69 | 0.56 | 0.98 | 0.85 | 0.52 | 0.48 | 0.58 | 0.99 | 0.78 | 0.44 | 0.56 | 0.57 | 1.00 | 0.81 |
|  | 4 min | 1.00 | 0.00 | 0.94 | 0.00 | 0.94 | 0.47 | 0.53 | 0.71 | 0.98 | 0.85 | 1.00 | 0.00 | 0.91 | 0.00 | 0.91 |
|  | 1 hr | 1.00 | 0.00 | 0.96 | 0.00 | 0.96 | 1.00 | 0.00 | 0.94 | 0.00 | 0.94 | 1.00 | 0.00 | 0.97 | 0.00 | 0.97 |
| 2_11 | 3 sec | 0.44 | 0.56 | 0.28 | 0.93 | 0.64 | 0.56 | 0.44 | 0.29 | 0.90 | 0.56 | 0.54 | 0.46 | 0.26 | 0.89 | 0.56 |
|  | 15 sec | 0.38 | 0.62 | 0.40 | 0.97 | 0.75 | 0.51 | 0.49 | 0.40 | 0.96 | 0.68 | 0.43 | 0.57 | 0.36 | 0.94 | 0.69 |
|  | 1 min | 0.27 | 0.73 | 0.49 | 0.95 | 0.83 | 0.50 | 0.50 | 0.52 | 0.97 | 0.75 | 0.33 | 0.67 | 0.45 | 0.94 | 0.78 |
|  | 4 min | 1.00 | 0.00 | 0.95 | 0.00 | 0.95 | 0.40 | 0.60 | 0.64 | 0.98 | 0.84 | 0.26 | 0.74 | 0.63 | 0.96 | 0.88 |
|  | 1 hr | 1.00 | 0.00 | 0.97 | 0.00 | 0.97 | 1.00 | 0.00 | 0.94 | 0.00 | 0.94 | 1.00 | 0.00 | 0.96 | 0.00 | 0.96 |
| 11_28 | 3 sec |  |  |  |  | 0.65 |  |  |  |  | 0.52 |  |  |  |  | 0.59 |
|  | 15 sec |  |  |  |  | 0.87 |  |  |  |  | 0.75 |  |  |  |  | 0.83 |
|  | 1 min |  |  |  |  | 0.93 |  |  |  |  | 0.85 |  |  |  |  | 0.90 |
|  | 4 min |  |  |  |  | 0.94 |  |  |  |  | 0.90 |  |  |  |  | 0.92 |
|  | 1 hr |  |  |  |  | 0.92 |  |  |  |  | 0.92 |  |  |  |  | 0.91 |
| 12_28 | 3 sec |  |  |  |  | 0.61 |  |  |  |  | 0.48 |  |  |  |  | 0.55 |
|  | 15 sec |  |  |  |  | 0.85 |  |  |  |  | 0.71 |  |  |  |  | 0.81 |
|  | 1 min |  |  |  |  | 0.91 |  |  |  |  | 0.83 |  |  |  |  | 0.88 |
|  | 4 min |  |  |  |  | 0.92 |  |  |  |  | 0.88 |  |  |  |  | 0.90 |
|  | 1 hr |  |  |  |  | 0.91 |  |  |  |  | 0.89 |  |  |  |  | 0.89 |
| 29_41 | 3 sec | 0.17 | 0.83 | 0.45 | 0.83 | 0.77 | 0.40 | 0.60 | 0.15 | 0.54 | 0.38 | 0.23 | 0.77 | 0.41 | 0.77 | 0.69 |
|  | 15 sec | 1.00 | 0.00 | 0.97 | 0.00 | 0.97 | 0.44 | 0.56 | 0.36 | 0.84 | 0.63 | 1.00 | 0.00 | 0.94 | 0.00 | 0.94 |
|  | 1 min | 1.00 | 0.00 | 0.98 | 0.00 | 0.98 | 0.21 | 0.79 | 0.39 | 0.90 | 0.79 | 1.00 | 0.00 | 0.96 | 0.00 | 0.96 |
|  | 4 min | 1.00 | 0.00 | 0.98 | 0.00 | 0.98 | 0.09 | 0.91 | 0.41 | 0.94 | 0.89 | 1.00 | 0.00 | 0.97 | 0.00 | 0.97 |
|  | 1 hr | 1.00 | 0.00 | 0.97 | 0.00 | 0.97 | 1.00 | 0.00 | 0.95 | 0.00 | 0.95 | 1.00 | 0.00 | 0.96 | 0.00 | 0.96 |
| 29_43 | 3 sec |  |  |  |  |  |  |  |  |  |  | 0.22 | 0.78 | 0.45 | 0.80 | 0.72 |
|  | 15 sec |  |  |  |  |  |  |  |  |  |  | 1.00 | 0.00 | 0.95 | 0.00 | 0.95 |
|  | 1 min |  |  |  |  |  |  |  |  |  |  | 1.00 | 0.00 | 0.97 | 0.00 | 0.97 |
|  | 4 min |  |  |  |  |  |  |  |  |  |  | 1.00 | 0.00 | 0.97 | 0.00 | 0.97 |
|  | 1 hr |  |  |  |  |  |  |  |  |  |  | 1.00 | 0.00 | 0.96 | 0.00 | 0.96 |
| 33_41 | 3 sec |  |  |  |  | 0.74 |  |  |  |  |  |  |  |  |  | 0.66 |
|  | 15 sec |  |  |  |  | 0.96 |  |  |  |  |  |  |  |  |  | 0.93 |
|  | 1 min |  |  |  |  | 0.96 |  |  |  |  |  |  |  |  |  | 0.96 |
|  | 4 min |  |  |  |  | 1.00 |  |  |  |  |  |  |  |  |  | 0.96 |
|  | 1 hr |  |  |  |  | 0.98 |  |  |  |  |  |  |  |  |  | 0.95 |
|  |  | Population |  | Deuteration |  | Avg | Population |  | Deuteration |  | Avg | Population |  | Deuteration |  | Avg |
|  |  | 1 | 2 | 1 | 2 |  | 1 | 2 | 1 | 2 |  | 1 | 2 | 1 | 2 |  |
| 42_62 | 3 sec | 0.15 | 0.85 | 0.49 | 0.83 | 0.78 | 0.32 | 0.68 | 0.32 | 0.71 | 0.59 | 0.22 | 0.78 | 0.48 | 0.79 | 0.72 |
|  | 15 sec | 0.05 | 0.95 | 0.46 | 0.97 | 0.95 | 0.20 | 0.80 | 0.45 | 0.89 | 0.80 | 1.00 |  | 0.94 | 0.00 | 0.94 |
|  | 1 min | 1.00 | 0.00 | 0.98 | 0.00 | 0.98 | 1.00 | 0.00 | 0.94 | 0.00 | 0.94 | 1.00 |  | 0.97 | 0.00 | 0.97 |
|  | 4 min | 1.00 | 0.00 | 0.99 | 0.00 | 0.99 | 1.00 | 0.00 | 0.98 | 0.00 | 0.98 | 1.00 |  | 0.98 | 0.00 | 0.98 |
|  | 1 hr | 1.00 | 0.00 | 0.98 | 0.00 | 0.98 | 1.00 | 0.00 | 0.98 | 0.00 | 0.98 | 1.00 |  | 0.97 | 0.00 | 0.97 |
| 54_62 | 3 sec | 0.27 | 0.73 | 0.53 | 0.84 | 0.76 | 0.31 | 0.69 | 0.36 | 0.76 | 0.64 |  |  |  |  | 0.70 |
|  | 15 sec | 1.00 | 0.00 | 0.97 | 0.00 | 0.97 | 1.00 | 0.00 | 0.88 | 0.00 | 0.88 |  |  |  |  | 0.94 |
|  | 1 min | 1.00 | 0.00 | 0.98 | 0.00 | 0.98 | 1.00 | 0.00 | 0.95 | 0.00 | 0.95 |  |  |  |  | 0.99 |
|  | 4 min | 1.00 | 0.00 | 0.99 | 0.00 | 0.99 | 1.00 | 0.00 | 0.98 | 0.00 | 0.98 |  |  |  |  | 1.00 |
|  | 1 hr | 1.00 | 0.00 | 0.99 | 0.00 | 0.99 | 1.00 | 0.00 | 0.99 | 0.00 | 0.99 |  |  |  |  | 0.99 |
| 55_62 | 3 sec |  |  |  |  | 0.78 |  |  |  |  | 0.71 |  |  |  |  | 0.72 |
|  | 15 sec |  |  |  |  | 0.96 |  |  |  |  | 0.91 |  |  |  |  | 0.94 |
|  | 1 min |  |  |  |  | 0.99 |  |  |  |  | 0.98 |  |  |  |  | 0.98 |
|  | 4 min |  |  |  |  | 0.99 |  |  |  |  | 1.00 |  |  |  |  | 0.98 |
|  | 1 hr |  |  |  |  | 0.98 |  |  |  |  | 0.99 |  |  |  |  | 0.98 |
| 55_76 | 3 sec |  |  |  |  | 0.95 |  |  |  |  | 0.89 |  |  |  |  | - |
|  | 15 sec |  |  |  |  | 1.00 |  |  |  |  | 0.96 |  |  |  |  | 0.96 |
|  | 1 min |  |  |  |  | 1.00 |  |  |  |  | 0.97 |  |  |  |  | 0.98 |
|  | 4 min |  |  |  |  | 1.01 |  |  |  |  | 0.99 |  |  |  |  | 0.99 |
|  | 1 hr |  |  |  |  | 1.00 |  |  |  |  | 0.99 |  |  |  |  | 0.98 |
| 63_73 | 3 sec |  |  |  |  | 0.97 |  |  |  |  | 0.94 |  |  |  |  | 0.96 |
|  | 15 sec |  |  |  |  | 0.99 |  |  |  |  | 0.98 |  |  |  |  | 0.99 |
|  | 1 min |  |  |  |  | 0.98 |  |  |  |  | 0.99 |  |  |  |  | 1.00 |
|  | 4 min |  |  |  |  | 0.99 |  |  |  |  | 1.00 |  |  |  |  | 1.01 |
|  | 1 hr |  |  |  |  | 0.98 |  |  |  |  | 1.00 |  |  |  |  | 1.00 |
| 63_76 | 3 sec |  |  |  |  | 0.98 |  |  |  |  | 0.95 |  |  |  |  | 0.97 |
|  | 15 sec |  |  |  |  | 0.99 |  |  |  |  | 0.98 |  |  |  |  | 0.98 |
|  | 1 min |  |  |  |  | 0.99 |  |  |  |  | 0.98 |  |  |  |  | 0.99 |
|  | 4 min |  |  |  |  | 0.98 |  |  |  |  | 0.99 |  |  |  |  | 0.99 |
|  | 1 hr |  |  |  |  | 0.98 |  |  |  |  | 0.99 |  |  |  |  | 0.98 |
| 63_84 | 3 sec |  |  |  |  | 0.98 |  |  |  |  |  |  |  |  |  |  |
|  | 15 sec |  |  |  |  | 0.98 |  |  |  |  |  |  |  |  |  |  |
|  | 1 min |  |  |  |  | 0.98 |  |  |  |  |  |  |  |  |  |  |
|  | 4 min |  |  |  |  | 0.98 |  |  |  |  |  |  |  |  |  |  |
|  | 1 hr |  |  |  |  | 0.97 |  |  |  |  |  |  |  |  |  |  |
| 63_99 | 3 sec |  |  |  |  | 0.80 | 0.24 | 0.76 | 0.47 | 0.77 | 0.70 |  |  |  |  |  |
|  | 15 sec |  |  |  |  | 0.83 | 0.23 | 0.77 | 0.57 | 0.80 | 0.75 |  |  |  |  |  |
|  | 1 min |  |  |  |  | 0.84 | 0.32 | 0.68 | 0.65 | 0.83 | 0.77 |  |  |  |  |  |
|  | 4 min |  |  |  |  | 0.88 | 0.27 | 0.73 | 0.69 | 0.87 | 0.82 |  |  |  |  |  |
|  | 1 hr |  |  |  |  | 0.90 | 1.00 | 0.00 | 0.88 | 0.00 | 0.88 |  |  |  |  |  |
| 77_84 | 3 sec |  |  |  |  | 0.98 |  |  |  |  | 0.94 |  |  |  |  |  |
|  | 15 sec |  |  |  |  | 0.98 |  |  |  |  | 0.96 |  |  |  |  |  |
|  | 1 min |  |  |  |  | 0.97 |  |  |  |  | 0.96 |  |  |  |  |  |
|  | 4 min |  |  |  |  | 0.97 |  |  |  |  | 0.98 |  |  |  |  |  |
|  | 1 hr |  |  |  |  | 0.96 |  |  |  |  | 0.97 |  |  |  |  |  |
| 77_87 | 3 sec |  |  |  |  | 0.97 | 0.21 | 0.79 | 0.44 | 0.94 | 0.84 |  |  |  |  | 0.97 |
|  | 15 sec |  |  |  |  | 0.97 | 0.28 | 0.72 | 0.63 | 0.96 | 0.87 |  |  |  |  | 0.96 |
|  | 1 min |  |  |  |  | 0.97 | 0.24 | 0.76 | 0.69 | 0.96 | 0.90 |  |  |  |  | 0.97 |
|  | 4 min |  |  |  |  | 0.97 | 0.29 | 0.71 | 0.76 | 0.99 | 0.92 |  |  |  |  | 0.98 |
|  | 1 hr |  |  |  |  | 0.96 | 1.00 | 0.00 | 0.96 | 0.00 | 0.96 |  |  |  |  | 0.96 |
| 77_99 | 3 sec |  |  |  |  | 0.69 | 0.29 | 0.71 | 0.32 | 0.67 | 0.57 |  |  |  |  | 0.66 |
|  | 15 sec |  |  |  |  | 0.72 | 0.26 | 0.74 | 0.38 | 0.69 | 0.61 |  |  |  |  | 0.69 |
|  | 1 min |  |  |  |  | 0.75 | 0.18 | 0.82 | 0.40 | 0.71 | 0.66 |  |  |  |  | 0.73 |
|  | 4 min |  |  |  |  | 0.79 | 0.15 | 0.85 | 0.46 | 0.77 | 0.72 |  |  |  |  | 0.79 |
|  | 1 hr |  |  |  |  | 0.84 | 1.00 | 0.00 | 0.81 | 0.00 | 0.81 |  |  |  |  | 0.83 |
|  |  | Population |  | Deuteration |  | Avg | Population |  | Deuteration |  | Avg | Population |  | Deuteration |  | Avg |
|  |  | 1 | 2 | 1 | 2 |  | 1 | 2 | 1 | 2 |  | 1 | 2 | 1 | 2 |  |
| 87_99 | 3 sec |  |  |  |  | 0.33 |  |  |  |  | 0.28 |  |  |  |  |  |
|  | 15 sec |  |  |  |  | 0.38 |  |  |  |  | 0.32 |  |  |  |  |  |
|  | 1 min |  |  |  |  | 0.45 |  |  |  |  | 0.40 |  |  |  |  |  |
|  | 4 min |  |  |  |  | 0.57 |  |  |  |  | 0.50 |  |  |  |  |  |
|  | 1 hr |  |  |  |  | 0.68 |  |  |  |  | 0.63 |  |  |  |  |  |
| 88_95 | 3 sec |  |  |  |  | 0.59 |  |  |  |  | 0.53 |  |  |  |  | 0.58 |
|  | 15 sec |  |  |  |  | 0.62 |  |  |  |  | 0.56 |  |  |  |  | 0.61 |
|  | 1 min |  |  |  |  | 0.64 |  |  |  |  | 0.61 |  |  |  |  | 0.63 |
|  | 4 min |  |  |  |  | 0.68 |  |  |  |  | 0.66 |  |  |  |  | 0.69 |
|  | 1 hr |  |  |  |  | 0.83 |  |  |  |  | 0.78 |  |  |  |  | 0.82 |
| 88_99 | 3 sec |  |  |  |  | 0.25 |  |  |  |  | 0.23 |  |  |  |  | 0.25 |
|  | 15 sec |  |  |  |  | 0.31 |  |  |  |  | 0.28 |  |  |  |  | 0.31 |
|  | 1 min |  |  |  |  | 0.38 |  |  |  |  | 0.36 |  |  |  |  | 0.40 |
|  | 4 min |  |  |  |  | 0.50 |  |  |  |  | 0.46 |  |  |  |  | 0.52 |
|  | 1 hr |  |  |  |  | 0.64 |  |  |  |  | 0.59 |  |  |  |  | 0.62 |
| 100_109 | 3 sec |  |  |  |  | 0.11 |  |  |  |  | 0.09 |  |  |  |  | 0.11 |
|  | 15 sec |  |  |  |  | 0.21 |  |  |  |  | 0.15 |  |  |  |  | 0.19 |
|  | 1 min |  |  |  |  | 0.29 |  |  |  |  | 0.21 |  |  |  |  | 0.27 |
|  | 4 min |  |  |  |  | 0.42 |  |  |  |  | 0.30 |  |  |  |  | 0.40 |
|  | 1 hr |  |  |  |  | 0.68 |  |  |  |  | 0.56 |  |  |  |  | 0.68 |
| 110_132 | 3 sec |  |  |  |  | 0.28 |  |  |  |  | 0.26 |  |  |  |  | 0.27 |
|  | 15 sec |  |  |  |  | 0.33 |  |  |  |  | 0.29 |  |  |  |  | 0.32 |
|  | 1 min |  |  |  |  | 0.38 |  |  |  |  | 0.33 |  |  |  |  | 0.36 |
|  | 4 min |  |  |  |  | 0.42 |  |  |  |  | 0.38 |  |  |  |  | 0.41 |
|  | 1 hr |  |  |  |  | 0.52 |  |  |  |  | 0.50 |  |  |  |  | 0.52 |
| 110_137 | 3 sec |  |  |  |  | 0.26 |  |  |  |  | 0.25 |  |  |  |  | 0.25 |
|  | 15 sec |  |  |  |  | 0.31 |  |  |  |  | 0.28 |  |  |  |  | 0.30 |
|  | 1 min |  |  |  |  | 0.35 |  |  |  |  | 0.31 |  |  |  |  | 0.34 |
|  | 4 min |  |  |  |  | 0.40 |  |  |  |  | 0.36 |  |  |  |  | 0.40 |
|  | 1 hr |  |  |  |  | 0.57 |  |  |  |  | 0.51 |  |  |  |  | 0.57 |
| 133_149 | 3 sec |  |  |  |  | 0.23 |  |  |  |  | 0.22 |  |  |  |  | 0.22 |
|  | 15 sec |  |  |  |  | 0.27 |  |  |  |  | 0.26 |  |  |  |  | 0.25 |
|  | 1 min |  |  |  |  | 0.30 |  |  |  |  | 0.28 |  |  |  |  | 0.28 |
|  | 4 min |  |  |  |  | 0.35 |  |  |  |  | 0.33 |  |  |  |  | 0.33 |
|  | 1 hr |  |  |  |  | 0.52 |  |  |  |  | 0.43 |  |  |  |  | 0.50 |
| 138_149 | 3 sec |  |  |  |  | 0.31 |  |  |  |  | 0.30 |  |  |  |  | 0.30 |
|  | 15 sec |  |  |  |  | 0.36 |  |  |  |  | 0.35 |  |  |  |  | 0.36 |
|  | 1 min |  |  |  |  | 0.41 |  |  |  |  | 0.39 |  |  |  |  | 0.39 |
|  | 4 min |  |  |  |  | 0.49 |  |  |  |  | 0.46 |  |  |  |  | 0.47 |
|  | 1 hr |  |  |  |  | 0.65 |  |  |  |  | 0.58 |  |  |  |  | 0.63 |
| 138_163 | 3 sec |  |  |  |  | 0.26 |  |  |  |  | 0.24 |  |  |  |  | 0.24 |
|  | 15 sec |  |  |  |  | 0.33 |  |  |  |  | 0.31 |  |  |  |  | 0.31 |
|  | 1 min |  |  |  |  | 0.40 |  |  |  |  | 0.38 |  |  |  |  | 0.38 |
|  | 4 min |  |  |  |  | 0.50 |  |  |  |  | 0.46 |  |  |  |  | 0.48 |
|  | 1 hr |  |  |  |  | 0.67 |  |  |  |  | 0.62 |  |  |  |  | 0.65 |
| 150_163 | 3 sec |  |  |  |  | 0.16 |  |  |  |  | 0.14 |  |  |  |  | 0.15 |
|  | 15 sec |  |  |  |  | 0.23 |  |  |  |  | 0.21 |  |  |  |  | 0.21 |
|  | 1 min |  |  |  |  | 0.32 |  |  |  |  | 0.29 |  |  |  |  | 0.30 |
|  | 4 min |  |  |  |  | 0.43 |  |  |  |  | 0.38 |  |  |  |  | 0.41 |
|  | 1 hr |  |  |  |  | 0.63 |  |  |  |  | 0.58 |  |  |  |  | 0.62 |
| 150_164 | 3 sec |  |  |  |  | 0.12 |  |  |  |  |  |  |  |  |  | 0.11 |
|  | 15 sec |  |  |  |  | 0.19 |  |  |  |  |  |  |  |  |  | 0.17 |
|  | 1 min |  |  |  |  | 0.27 |  |  |  |  |  |  |  |  |  | 0.25 |
|  | 4 min |  |  |  |  | 0.37 |  |  |  |  |  |  |  |  |  | 0.35 |
|  | 1 hr |  |  |  |  | 0.57 |  |  |  |  |  |  |  |  |  | 0.56 |
|  |  | Population |  | Deuteration |  | Avg | Population |  | Deuteration |  | Avg | Population |  | Deuteration |  | Avg |
|  |  | 1 | 2 | 1 | 2 |  | 1 | 2 | 1 | 2 |  | 1 | 2 | 1 | 2 |  |
| 164_172 | 3 sec |  |  |  |  | 0.41 |  |  |  |  | 0.41 |  |  |  |  | 0.41 |
|  | 15 sec |  |  |  |  | 0.42 |  |  |  |  | 0.41 |  |  |  |  | 0.42 |
|  | 1 min |  |  |  |  | 0.44 |  |  |  |  | 0.43 |  |  |  |  | 0.45 |
|  | 4 min |  |  |  |  | 0.53 |  |  |  |  | 0.50 |  |  |  |  | 0.55 |
|  | 1 hr |  |  |  |  | 0.62 |  |  |  |  | 0.61 |  |  |  |  | 0.62 |
| 164_178 | 3 sec |  |  |  |  | 0.70 |  |  |  |  | 0.69 |  |  |  |  | 0.68 |
|  | 15 sec |  |  |  |  | 0.70 |  |  |  |  | 0.69 |  |  |  |  | 0.68 |
|  | 1 min |  |  |  |  | 0.71 |  |  |  |  | 0.70 |  |  |  |  | 0.70 |
|  | 4 min |  |  |  |  | 0.76 |  |  |  |  | 0.74 |  |  |  |  | 0.75 |
|  | 1 hr |  |  |  |  | 0.79 |  |  |  |  | 0.78 |  |  |  |  | 0.78 |
| 165_178 | 3 sec |  |  |  |  | 0.76 |  |  |  |  | 0.75 |  |  |  |  |  |
|  | 15 sec |  |  |  |  | 0.76 |  |  |  |  | 0.76 |  |  |  |  |  |
|  | 1 min |  |  |  |  | 0.77 |  |  |  |  | 0.77 |  |  |  |  |  |
|  | 4 min |  |  |  |  | 0.83 |  |  |  |  | 0.81 |  |  |  |  |  |
|  | 1 hr |  |  |  |  | 0.86 |  |  |  |  | 0.86 |  |  |  |  |  |
| 179_185 | 3 sec | 0.20 | 0.80 | 0.36 | 0.93 | 0.81 | 0.33 | 0.67 | 0.36 | 0.92 | 0.74 | 0.15 | 0.85 | 0.30 | 0.83 | 0.75 |
|  | 15 sec | 1.00 |  | 0.97 | 0.00 | 0.97 | 0.13 | 0.87 | 0.45 | 0.98 | 0.91 | 1.00 |  | 0.94 | 0.00 | 0.94 |
|  | 1 min | 1.00 |  | 0.94 | 0.00 | 0.94 | 1.00 |  | 1.01 | 0.00 | 1.01 | 1.00 |  | 0.95 | 0.00 | 0.95 |
|  | 4 min | 1.00 |  | 0.95 | 0.00 | 0.95 | 1.00 |  | 1.04 | 0.00 | 1.04 | 1.00 |  | 0.96 | 0.00 | 0.96 |
|  | 1 hr | 1.00 |  | 0.98 | 0.00 | 0.98 | 1.00 |  | 0.99 | 0.00 | 0.99 | 1.00 |  | 0.98 | 0.00 | 0.98 |
| 179_205 | 3 sec |  |  |  |  | 0.93 |  |  |  |  | 0.92 |  |  |  |  | 0.91 |
|  | 15 sec |  |  |  |  | 0.98 |  |  |  |  | 0.96 |  |  |  |  | 0.96 |
|  | 1 min |  |  |  |  | 0.98 |  |  |  |  | 0.97 |  |  |  |  | 0.97 |
|  | 4 min |  |  |  |  | 0.99 |  |  |  |  | 0.98 |  |  |  |  | 0.98 |
|  | 1 hr |  |  |  |  | 0.98 |  |  |  |  | 0.98 |  |  |  |  | 0.97 |
| 186_201 | 3 sec |  |  |  |  | 0.97 |  |  |  |  | 0.97 |  |  |  |  | 0.97 |
|  | 15 sec |  |  |  |  | 0.96 |  |  |  |  | 0.96 |  |  |  |  | 0.96 |
|  | 1 min |  |  |  |  | 0.96 |  |  |  |  | 0.96 |  |  |  |  | 0.97 |
|  | 4 min |  |  |  |  | 0.96 |  |  |  |  | 0.97 |  |  |  |  | 0.97 |
|  | 1 hr |  |  |  |  | 0.95 |  |  |  |  | 0.96 |  |  |  |  | 0.96 |
| 186_205 | 3 sec |  |  |  |  | 0.95 |  |  |  |  | 0.95 |  |  |  |  | 0.94 |
|  | 15 sec |  |  |  |  | 0.97 |  |  |  |  | 0.97 |  |  |  |  | 0.96 |
|  | 1 min |  |  |  |  | 0.98 |  |  |  |  | 0.97 |  |  |  |  | 0.98 |
|  | 4 min |  |  |  |  | 0.97 |  |  |  |  | 0.98 |  |  |  |  | 0.98 |
|  | 1 hr |  |  |  |  | 0.97 |  |  |  |  | 0.98 |  |  |  |  | 0.97 |
| 187_205 | 3 sec |  |  |  |  | 0.95 |  |  |  |  | 0.95 |  |  |  |  | 0.94 |
|  | 15 sec |  |  |  |  | 0.97 |  |  |  |  | 0.97 |  |  |  |  | 0.97 |
|  | 1 min |  |  |  |  | 0.97 |  |  |  |  | 0.97 |  |  |  |  | 0.98 |
|  | 4 min |  |  |  |  | 0.97 |  |  |  |  | 0.98 |  |  |  |  | 0.98 |
|  | 1 hr |  |  |  |  | 0.97 |  |  |  |  | 0.98 |  |  |  |  | 0.97 |

